# Music Scaffolds Visual Statistical Sequence Learning Through Network-Level Reorganization in the Brain

**DOI:** 10.1101/2025.08.05.668768

**Authors:** Yiren Ren, Vishwadeep Ahluwalia, Claire Arthur, Thackery Brown

**Affiliations:** School of Psychology, College of Sciences, Georgia Institute of Technology; Center for Advanced Brain Imaging, Georgia Institute of Technology/Georgia State University; School of Music, College of Design, Georgia Institute of Technology

## Abstract

Statistical learning—the ability to extract patterns from noisy continuous experiences—is fundamental to human cognition. Yet, how contextual factors shape this process remains poorly understood. Music is an important example of such contextual factors, because it is ubiquitous in human experience and provides a rich temporally-structured stimulus that can co-occur with other learning processes. Here we demonstrate that pairing music fundamentally enhances visual statistical learning, and this is correlated with systematic reorganization of large-scale brain networks. Using fMRI and a novel probabilistic sequence learning paradigm, we show that familiar melodies significantly improved participants’ ability to segment continuous visual streams into events and learn sequential relationships. Neuroimaging analyses revealed that the presence of music fundamentally altered the neural network organization that coordinates learning mechanisms: while sequence learning in silence engaged frontal-parietal networks associated with explicit pattern extraction, providing musical temporal structure as a context shifted learning toward MTL-vmPFC circuits recently implicated in schema-guided memory processing. Machine learning analyses confirmed these architectural differences, with the music condition achieving optimal neural prediction of behavioral performance through distributed connectivity patterns while control condition relied on concentrated processing. Our findings support a Cross-Modal Temporal Scaffolding Theory, demonstrating that structured temporal context signals from one modality (here, music) can create more efficient neural states for sequence processing in another through dual mechanisms: enhanced memory integration through schema-guided learning and reduced demands on explicit control resources. These results identify network-level principles for optimizing statistical learning, with broad implications for understanding how environmental context shapes human learning capacity.

## Introduction

In human daily life, our brain constantly receives continuous and stochastic inputs. The ability to segment these noisy inputs into meaningful groups and extract regular sequential relationships between events or items is fundamental to numerous aspects of human cognition and behavior, underpinning essential skills such as language comprehension, navigation, motor skill development, and autobiographical memory recall. Successfully navigating our environment conceptually and physically requires understanding temporal relationships and statistical regularities amid variable input - a core adaptive mechanism that allows humans to make predictions and organize experiences into meaningful episodes (Kurby & Zacks, 2008; Sherman et al., 2020; Tulving, 2002).

Statistical learning paradigms have emerged as a powerful tool for investigating how humans extract probabilistic regularities from variable input streams, enabling prediction and pattern recognition even when individual experiences are noisy or incomplete (Saffran et al., 1996; Turk-Browne et al., 2005). Unlike deterministic associative learning, which involves memorizing fixed stimulus-stimulus or stimulus-response relationships, statistical learning requires computing distributional properties across multiple exposures to identify underlying structure amid variability (Frost et al., 2015). This process is fundamental to real-world cognition, where perfect repetitions are rare and learners must instead discern reliable patterns from inconsistent input—such as extracting word boundaries from continuous speech or predicting environmental events from probabilistic cues. While early work conceptualized statistical learning as primarily implicit, mounting evidence suggests that both implicit and explicit knowledge acquisition can reveal statistical learning effects (Batterink et al., 2015; Franco et al., 2011), and can be supported by “declarative” memory circuitry in the brain (Covington et al., 2018; Schapiro et al., 2016, 2017; Schlichting et al., 2017). These mechanisms have been demonstrated across multiple domains and extend to complex real-world scenarios and hierarchical regularities (Aslin & Newport, 2012; Brady & Oliva, 2008; Fiser & Aslin, 2002; Loui et al., 2010; Schapiro et al., 2014). Recent comprehensive reviews have highlighted both the domain-general nature of statistical learning processes and the critical role of modality-specific constraints (Conway, 2020; Frost et al., 2015), with mounting evidence that these mechanisms operate across multiple domains while exhibiting systematic modality and stimulus specificity.

Critically, in real-world settings, statistical learning rarely occurs in isolation—we usually encode sequences and extract temporal patterns while multiple contextual factors are simultaneously present. Consider watching a film, where we process visual narrative sequences accompanied by a musical score; or learning a step-by-step procedure while background music plays; or driving a daily commute route while listening to the radio. A fundamental question emerges: does structured temporal information in one modality facilitate or hinder our ability to extract statistical regularities in another? While research has established that statistical learning can operate across multiple modalities simultaneously (Conway & Christiansen, 2006; Mitchel & Weiss, 2011; Seitz et al., 2007), a critical gap remains in understanding how structured temporal context from one modality specifically scaffolds pattern extraction in another. The auditory scaffolding hypothesis (Conway et al., 2009) proposed that auditory temporal structure may be particularly well-suited to support temporal processing across domains, but the neural mechanisms underlying such cross-modal temporal scaffolding remain largely unexplored.

### Why is music positioned to provide such benefits to statistical learning?

Literature indicates that context significantly modulates statistical and sequential learning in part by defining the temporal and spatial boundaries of events (Tulving, 1983), and creating natural parsing points in continuous experience (DuBrow & Davachi, 2013). Shared context facilitates the binding of sequential elements into coherent memory representations, while context changes create distinct separations between events in memory (Heusser et al., 2018; Pettijohn et al., 2016). Schema literatures have shown that when new information maps onto existing knowledge structures, learning becomes more efficient through enhanced communication between the prefrontal cortex and hippocampus (van Kesteren et al., 2012). If music’s temporal structure facilitates extracting the sequential relationships of stimuli in the environment despite variability in those stimuli across repetitions, it would provide compelling evidence that cross-modal scaffolding can enhance statistical learning. These properties—hierarchical temporal structure, schematic familiarity, and predictable temporal cues—position familiar music as an ideal candidate for investigating cross-modal temporal enhancement effects.

Our recent work has begun to systematically investigate when and how these musical properties influence visual sequence learning. Our initial studies demonstrated that familiar, structurally regular music enhances deterministic sequence learning compared to both monotonic auditory controls and unfamiliar musical pieces (Ren et al., 2024; Ren & Brown, 2025). These findings revealed that both familiarity and structural regularity are critical factors, with preliminary analyses indicating that monotonic auditory streams produce statistically equivalent effects to silence conditions. This systematic investigation established specific constraints for musical enhancement effects, demonstrating when musical scaffolding occurs. Having established their behavioral benefits empirically, the current study shifts focus from efficacy testing to mechanistic investigation – examining how these effects emerge through neural network reorganization. Rather than re-testing whether music helps sequence learning, we investigate the brain mechanisms underlying this empirically-validated phenomenon.

Based on these empirical foundations, we propose a Cross-Modal Temporal Scaffolding Theory (CMTS; ***Figure 1***) to explain when and how structured temporal information from one modality enhances learning in another. CMTS builds upon established theoretical frameworks including schema theory (van Kesteren et al., 2012), which suggests that new information is more efficiently encoded when it aligns with existing knowledge structures; complementary learning systems theory (McClelland et al., 2020), which proposes interactions between rapid and slow learning systems; and Baddeley’s working memory model (2020), which describes partial segregation of modalities that can mitigate interference effects.

**Figure 1.**
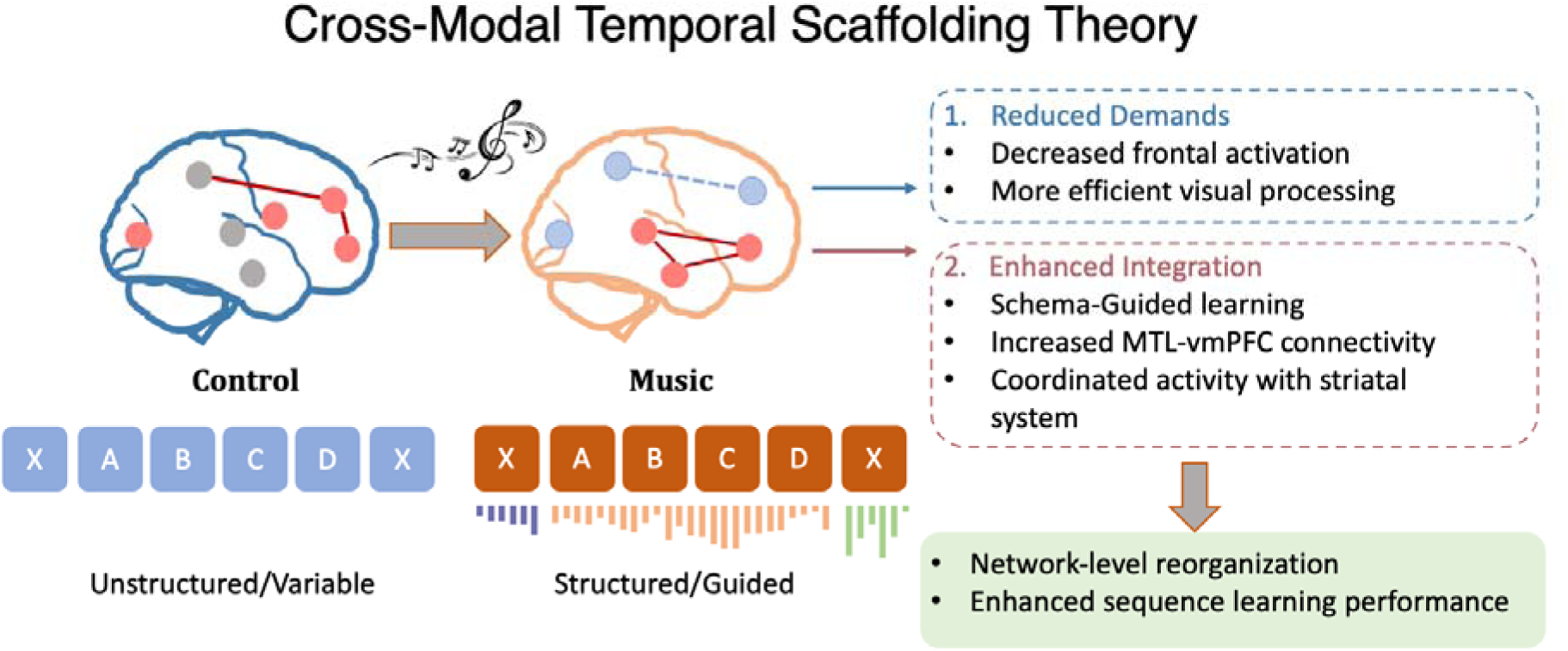
Cross-Modal Temporal Scaffolding Theory. *Musical context shifts neural processing from variable frontal-parietal networks (left) to consistent MTL-vmPFC-striatal circuits (right) through two mechanisms: (1) Reduced demands on explicit control resources, resulting in decreased frontal activation and more efficient processing; (2) Enhanced memory integration, characterized by increased MTL-vmPFC connectivity and schema-guided learning. This network reorganization leads to enhanced sequential learning performance and more predictable relationships between neural activity and behavioral outcomes*.

CMTS integrates these perspectives, proposing that temporal scaffolding operates through dual mechanisms: (1) enhanced memory integration via schema-guided learning, where familiar temporal structures facilitate encoding through established MTL-PFC circuits, and (2) reduced demands on explicit control resources, shifting processing from effortful frontal-parietal networks toward more efficient memory-based systems. Critically, this framework integrates convergent data (including the results in this report) showing that when appropriately structured, cross-modal temporal cues can create synergistic rather than competitive effects between modalities. Importantly, CMTS predicts that musical scaffolding should extend beyond simple boundary detection to enhance relational memory encoding itself. While any temporal cue might facilitate segmentation, structured musical schemas should enhance the encoding of sequential relationships between items within sequences through established memory integration pathways. This distinction allows CMTS to make specific predictions about both behavioral patterns (benefits extending beyond boundary detection to sequential order memory) and neural mechanisms (enhanced MTL-PFC connectivity reflecting memory integration rather than attention/segmentation networks alone).

Statistical learning engages coordinated activity across distributed brain networks including the medial temporal lobe (MTL) for detecting and binding sequential elements (Hsieh et al., 2014; Schapiro et al., 2014, 2016), the prefrontal cortex (PFC) for temporal information maintenance(Blumenfeld et al., 2011), and the striatum for rhythmic patterns processing and predications (Bornstein & Daw, 2012; Grahn & Rowe, 2009; Meck et al., 2008). These regions work with broader networks such as inferior frontal (Karuza et al., 2013; Petersson et al., 2012; Turk-Browne et al., 2009) and parietal cortices (Grabner et al., 2009; Karuza et al., 2013; Thomas et al., 2004), which integrate and extract probabilistic patterns for successful regularity learning. Recent neuroimaging investigations have further characterized these networks, revealing hierarchical sequence representations across cortex and hippocampus (Henin et al., 2021), frontoparietal networks that explain individual differences in statistical learning (Orpella et al., 2022), and meta-analytic evidence for consistent activation patterns during artificial grammar learning (Ramage et al., 2024). Building on this foundation, we investigated whether contextual factors can systematically reorganize these established learning networks.

The CMTS framework generates specific neural predictions for how musical context should modulate these networks and reduce individual variability. First, music as a temporal context signal should drive more efficient network configurations by providing ‘temporal schemas’—familiar musical structures should facilitate encoding through enhanced MTL-PFC connectivity, replicating our previous finding (Ren & Brown, 2025), similar to how existing knowledge structures facilitate new learning (van Kesteren et al., 2012). This schema-guided processing should strengthen connections between the hippocampus, vmPFC, and striatum— regions critical for integrating temporal context with sequential memories (Grahn & Rowe, 2009; Preston & Eichenbaum, 2013). Second, this network reorganization should reduce demands on effortful frontal-parietal pathways, particularly IFG-angular gyrus circuits typically engaged during explicit pattern extraction under uncertainty (Grabner et al., 2009; Kim et al., 2022). Finally, this dual-mechanism model should manifest in more consistent neural states across participants when musical structure provides temporal organization, reflecting optimized engagement of memory-based learning circuits rather than idiosyncratic control processes. These predictions are grounded in CMTS constraints established through our systematic research: musical temporal scaffolding requires both familiarity (to activate existing schemas) and structural regularity (to provide predictable temporal organization). Statistical learning paradigms provide an optimal test of this framework because extracting probabilistic patterns from noisy input streams places greater demands on cognitive resources than deterministic sequence learning, making the potential benefits of temporal scaffolding more pronounced.

The present study tests these predictions using whole-brain fMRI during a visual statistical learning task where participants extracted probabilistic sequential patterns from continuous image streams. Using univariate activation, region-of-interest, and functional connectivity analyses, we characterized how musical context influences brain network organization during statistical learning, testing whether cross-modal temporal scaffolding drives the predicted shift from variable frontal-parietal engagement toward more consistent MTL-PFC-striatal connectivity.

## Method

### Participants

Eighty-nine healthy participants (42 males and 47 females) were recruited from the Georgia Institute of Technology’s volunteer pool. Participants’ ages range from 18 to 35, with an average of 20.57 years and standard deviation of 3.27 years old. Out of the 89 participants, 36 underwent fMRI scanning, while the remaining 53 participants completed the task with behavioral measures only. Among the 36 fMRI participants, 12 were males and 24 were females. These participants ranged from 18 years old to 35 years old (mean = 21.58, std = 4.1). However, one participant encountered technical issues during the MRI session, and stimuli timings were not encoded correctly. As a result, this participant was excluded from the MRI analysis, leaving the final fMRI sample size at 35. Sample size determination was guided by both behavioral and neuroimaging power considerations. For behavioral effects, power analysis using G*Power (Faul et al., 2009) based on preliminary data from the pilot study indicated that 73 participants would provide >80% power to detect medium-sized effects (d > 0.4) at α = 0.05. Our final behavioral sample of 89 participants exceeded this requirement. For the fMRI component, our sample size of 35 participants was determined based on successful detection of music-modulated sequence learning effects in our previous study (Ren and Brown, 2025) using a similar sample size. Prior to the study, all participants completed a self-report questionnaire to ensure they did not have any of the following conditions that could affect the study outcome: auditory impairments, visual impairments not corrected by glasses or contacts, fundamental music processing deficits (e.g., amusia or music agnosia), learning disabilities, attention disorders, or a history of neurological or psychiatric illnesses. Participants who volunteered for the fMRI scan were subject to further screening to exclude any magnetic resonance imaging contraindications such as metallic implants or major neurological disorders, following the Georgia Institute of Technology/Georgia State University Center for Advanced Brain Imaging’s scanning policy. All participants provided informed consent to the procedures approved by the Institutional Review Board of Georgia Institute of Technology.

### Materials

#### Visual Stimuli

Visual stimuli comprised real-life images selected from diverse databases, including the 360-color items dataset (Moreno-Martínez & Montoro, 2012), MIT indoors scenes (Quattoni & Torralba, 2009) and visual stimuli used and shared by our lab in prior studies (e.g. Brown et al., 2020). These images encompassed categories such as objects, animals, and scenes with the goal that memory results would better reflect what one might expect for the types of complex and often arbitrary associations experienced in the course of daily life. Each image was adjusted to a size of 300x300 pixels. A total of 72 distinct images were chosen to form 18 sequences, each composed of four images. This sequence length was chosen based on previous research (Ren et al., 2024) indicating that four items per sequence allow for sufficient learning within a two-hour session without reaching a ceiling effect. 12 sequences were assigned to the music condition and the remaining 6 to the control condition (see details in Procedure). This allocation was determined by this study being part of a broader research project investigating different types of music, which constitutes a separate study that will be reported in another paper. For the purposes of this paper, all music conditions are collapsed into a single ‘music present’ condition to address our primary research question about the general effects of musical context on sequence learning. The control sequences provide a baseline measure of sequence learning without musical accompaniment. The object, scene, and animal categories were balanced across the music and control conditions, as well as across sequences within each condition. However, the specific composition of each sequence varied to avoid introducing additional regularities that could influence learning. For example, while each sequence contained no more than one scene image, some sequences did not include any scenes. The assignment of specific images to sequences was randomized to control for potential biases. An additional set of 50 images, aligned with the sequence categories, served as lures during encoding, interspersed between sequences.

#### Music Stimuli

Music stimuli were selected from a recently published famous melodies stimulus set (Belfi & Kacirek, 2021) and popular classical music compositions in the literature, for a total of 12 melodies. Based on our prior study (Ren et al., 2024), music familiarity was an important factor for memory outcomes; to select these 12 melodies we first selected the top 20 melodies with highest ‘familiarity’ score that resonated with both young and old participants from Belfi and Kacirek, 2021. Additionally, we incorporated ten classical pieces frequently used in music cognition literature. We then put these 30 musical pieces into a pilot survey before running the experiment, asking an independent set of 20 people to rate the familiarity of each song. Questions included: 1) rate how fast you can recognize the music from not until the end to immediately; 2) if the music is paused at the fifth second, can you retrieve the melody and keep singing the rest of the song; 3) can you sing along the music when you listen to it either out loud or in your mind? 4) rate the overall familiarity from 0 indicating you don’t know the song to 10 indicating you can recall and sing this song out loud without any cue. Following this validation survey with 20 participants, we selected the 12 melodies with the highest familiarity score to be used in this study (participants in the experiment sample further validated their familiarity to be eligible; 2.3.1):

1. "Blue Danube"
2. "Deck the Halls"
3. "Eine Kleine"
4. "Für Elise"
5. "Harry Potter Theme Music"
6. "Itsy Bitsy Spider"
7. "Jingle Bells"
8. "Ode to Joy"
9. "Alla Turca" from Sonata No.11 in A major, K331 (Mozart)
10. "Pop Goes the Weasel"
11. "Rudolph the Red-Nosed Reindeer"
12. "Wedding March" from ‘Lohengrin’

These 12 melodies were paired with 12 visual sequences. To control relevant music parameters across stimuli, we used Logic Pro music composition software to extract the most recognizable melodic segments from each piece, rather than using entire compositions. Each excerpt was arranged for piano, standardizing the timbre across all stimuli. The primary melodic lines were set at 90 dB, with accompanying harmonies at 70 dB to ensure the melody remained prominent while maintaining consistent audio characteristics across all selections. To maintain the recognizability of each melody while ensuring consistent presentation duration, we selected the most identifiable segment from each piece and adjusted it to fit within an 8-second timeframe. This adjustment preserved each melody’s characteristic tempo (ranging from 160-240 BPM across selections) while standardizing the overall duration. We prioritized maintaining each melody’s natural rhythm and musical identity, as this familiarity was crucial for our experimental design. Each 8-second excerpt was then structured into four equal 2-second phrases, with a strong metrical accent at the beginning of each phrase to align precisely with the onset of each visual stimulus. This approach ensured consistent temporal alignment between musical phrases and visual stimuli while preserving the distinctive and recognizable qualities of each familiar melody.

For the lure images (non-member images of the sequences) inserted during the encoding of visual image streams, we used Mozart’s piano sonata No. 16 as background sound. This piece was chosen as it met several practical criteria: it has sufficient length for the lure presentation duration (144 seconds), maintains consistent musical style and instrumentation with the sequence-paired stimuli, and was not among the highly familiar melodies used in the main task to avoid confusion. This music was generated using Logic-Pro in the same way as other music stimuli, except for a longer duration (144 seconds).

The consistent music-sequence pairings in our design were chosen to optimize temporal scaffolding effects, informed by our broader systematic research program. Our task battery includes three complementary measures: boundary detection (segmentation accuracy), order recognition during encoding (retrieval practice), and complete sequence reconstruction (final test). This multi-measure approach enables assessment of whether benefits are limited to segmentation (consistent with boundary cueing) or extend to relational memory encoding (consistent with temporal scaffolding). Additionally, our neuroimaging analyses focus on both regional activation patterns and network connectivity to distinguish between attention/segmentation systems typically engaged by temporal cues versus memory integration circuits predicted by Cross-Modal Temporal Scaffolding Theory.

### Experiment Procedure

The experiment consisted of four main components: 1) pre-task screening (> 3 days before the task); 2) practice trials (5 minutes); 3) encoding phase with embedded retrieval practice (in scanner, around 2 hours); 4) final retrieval task (post-scan, 10 minutes)

#### Participant Pre-Task Screening

Prior to participating, all participants underwent an online pre-task screening questionnaire using Google Form at least three days in advance. They were required to listen to audio clips of the 12 pre-selected music stimuli and rate their familiarity on a scale of 1 to 5 with the questions "Are you familiar with the music?" and “Can you sing along with the melody without any problem?”. Only participants who answered all question with a score of 5 were eligible to participate in the task.

#### Experimental Procedure

The entire experiment was conducted using MATLAB Psychtoolbox 3 and comprised two distinct phases: encoding and retrieval. The encoding phase took place within the MRI scanner, where participants were exposed to continuous streams of images, some of which were accompanied by music. Participants were instructed to learn the sequential relationships among the images and to indicate their perception of event boundaries. Retrieval practice tasks were inserted within the encoding phase, and a subsequent sequence recall task was used after the scanning session. These retrieval tasks assessed participants’ memory for the learned sequences and individual items. The combination of encoding and retrieval phases allowed for a comprehensive examination of the effects of musical context on temporal order learning and memory.

##### Encoding Phase

The encoding phase took place within the MRI scanner. Participants were instructed that they would be presented with a continuous stream of images and their goal was to extract any regular patterns and learn the sequential relationship of images. To demonstrate their statistical learning of these patterns, they were instructed to perform a segmentation task during image presentation, pressing a button on an MRI-compatible response box whenever they sensed a boundary or breaking point of an event (group).Unknown to the participants, the encoding phase contained 18 repeating sequences embedded in a continuous stream of images, surrounded by lure stimuli that occurred stochastically. Among these, 12 sequences were paired with music, with each sequence consistently corresponding to the same music composition for one participant. The sequence-music pairings were randomly assigned at the start for each participant and remained constant throughout. The remaining six sequences were learned without any accompanying music, serving as a control condition.

While our previous studies used an active non-musical control (monotonic tone sequences) and demonstrated that musical structure provides benefits beyond mere rhythmic cues (Ren et al., 2024), the current study opted for a silence condition to examine how musical context affects learning compared to naturalistic baseline conditions. This choice was motivated by two considerations: First, our previous works (Ren et al., 2024; Ren & Brown, 2025) had already established music has advantages over simple rhythmic but non-musical cues, allowing us to focus here on understanding how musical structure alters neural mechanisms relative to typical learning scenarios in sequencing experiments. Second, visual sequential learning in everyday life often occurs without accompanying auditory input, making silence an ecologically valid control condition for understanding how musical context modifies natural learning processes.

Participants were explicitly instructed to focus solely on learning the visual sequences. They were told that the presence of music was incidental and that they would not be tested on any aspect of the auditory information. This instruction was given to prevent participants from trying to find explicit associations between the music and visual sequences or overthinking the experimental design.

During encoding, shown in ***Figure 2,*** participants viewed continuous streams of images. Each image was presented for 2 seconds. There was a fixed 2/3 probability that each sequence would be displayed in the correct fixed order, while there was a 1/3 probability that the sequence would be randomly shuffled. This specific probability was chosen based on a combination of pilot testing and practical considerations. Pilot studies were conducted with varying probabilities to identify a level of difficulty that was challenging enough to require statistical learning but not so difficult as to make learning impossible within the given time constraints. Moreover, the 2/3 probability allowed for a balanced design that could accommodate the desired number of trials within the 2-hour scanning session. In the presence of music, each musical excerpt was also 8 seconds long, corresponding to the duration of a single sequence. If the sequence was shuffled, the music was also correspondingly shuffled—each two seconds of the music paired with a specific picture within the sequence – for the present study, this ensured the syntactical/semantic structure of music was a maximally meaningful learning cue rather than the relationships between visual items and tonal items themselves being probabilistic (although this would be an interesting extension for future work). To prevent participants from simply segmenting events every eight seconds, various distracting images were interspersed between sequences, in both control and music conditions. Concurrently, a random section from Mozart’s piano sonata No.16 was played. This probabilistic presentation of sequences, along with the inclusion of lure images, was designed to create a statistical learning task where participants had to extract regularities from a noisy input stream. By contrast, a deterministic repetition of stimuli would not require the same level of statistical computation and would instead rely more on rote memorization. The current design aims to capture the complex and probabilistic nature of real-world learning scenarios, where perfect repetitions are rare, and learners must discern patterns amidst noise and variability.

**Figure 2.**
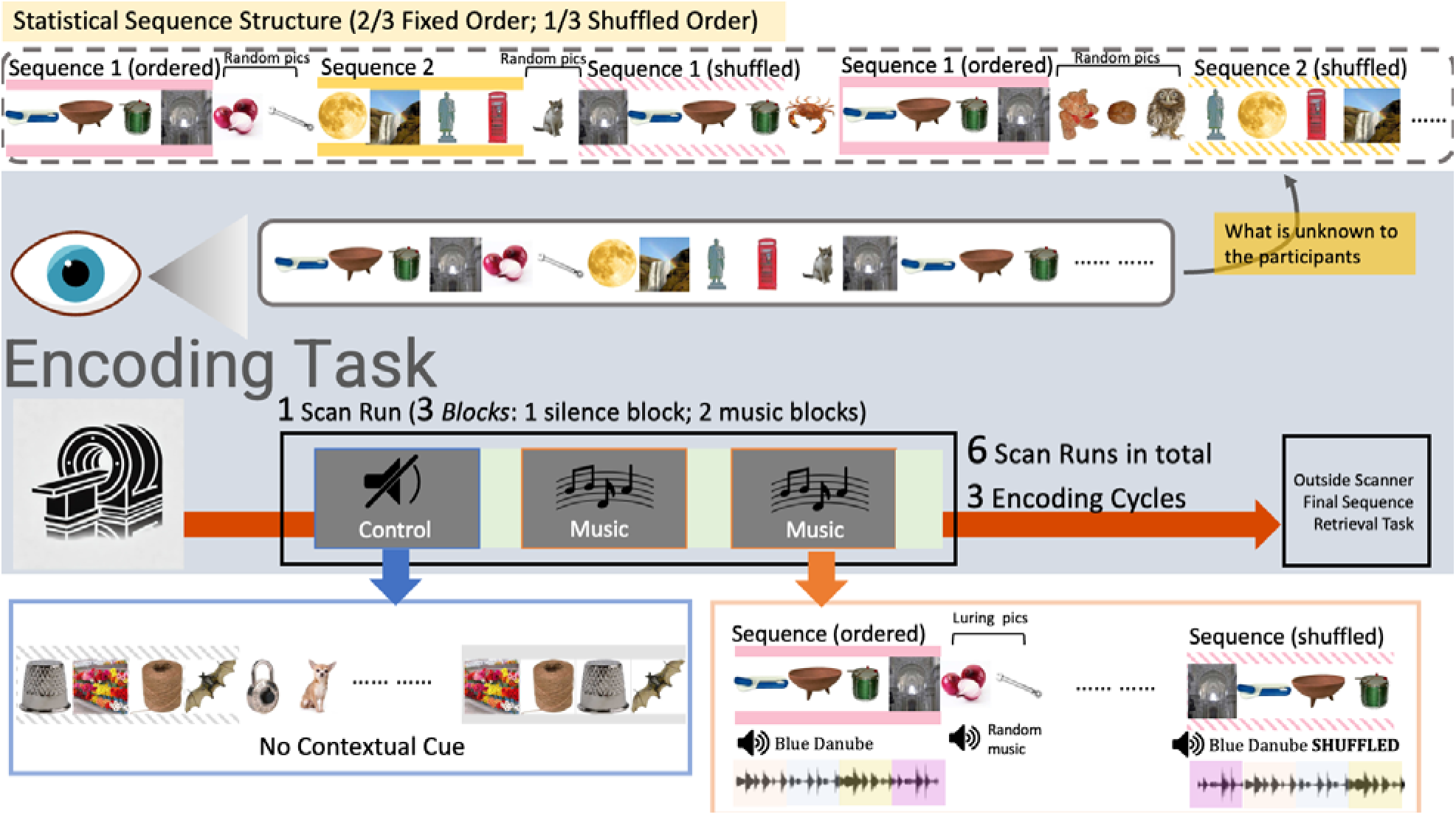
Experiment Procedure. *During encoding, participants viewed continuous streams of images inside an MRI scanner. (**Upper Panel**) What is unknown to the participants is that the visual sequences were probabilistically ordered (2/3 correct/fixed and 1/3 shuffled). Random numbers of random pictures were inserted between sequences to prevent segmenting sequences every four images. (**Center Panel**) Each scan run lasts around 10-12 minutes, consisting of 3 blocks (one in control condition and two in music condition). Participants were asked to 1) try to learn and extract the sequential relationships of visual images; 2) segment events by pressing a button when they perceived a boundary. Retrieval practice tasks followed each block, testing sequence order recall (see* Fig 3*). After fMRI scanning, a final retrieval task was given (see* Fig 3*). There were six scan runs in total, all sequences were learned in every two runs, making three encoding cycles in total for each sequence. (**Bottom Panel**) During control condition, participants viewed visual sequences in silence. During music condition, each sequence paired with a specific familiar melody. Importantly, each image in a sequence always paired with the same segment of the melody. Therefore, when the sequence was shuffled, the music audio would also be shuffled accordingly*.

The experiment was conducted over six scanning runs, interspersed with in-scanner rest periods and scanner restarts. Each RUN lasted approximately eleven minutes. This allowed participants to relax and rest which is critical for their performance due to the length of the scan (2 hours in total). Within each run, participants had three learning BLOCKs, divided by retrieval practice tasks. Among these three blocks, two were music condition and one was control condition. In each block, participants learned three sequences in total, with each sequence appearing six times and having a 1/3 chance of being shuffled. The order of the sequences and blocks was randomized. Participants completed one repetition of learning all 18 sequences in every two scanning runs, making it three repetitions of learning in total during the scan.

Below summarizes the main sequence presentation parameters:

- Probability of correct order: 2/3
- Probability of shuffled order: 1/3
- Number of repetitions per sequence: 6 times per block
- Number of blocks per scanning run: 3
- Number of scanning runs: 6
- Total repetitions of sequences learning across experiment: 3 (across all runs)

##### Retrieval Practice Tasks

Retrieval practice tasks were inserted after every three and half minutes of image presentation (after one block where three sequences were learned), resulting in three retrieval practice rounds per run. The inclusion of these tasks served two crucial purposes. First, based on early pilot studies, it was found that participants experienced difficulty maintaining focus and learning if the continuous stream of images was too long. Therefore, the decision was made to divide each run into three blocks, with retrieval practice tasks providing a mental break for participants. More importantly, while segmentation presses offered insight into participants’ learning in terms of their ability to segment and cluster visual items, the retrieval practice tasks provided a measure of their learning of the temporal order relationships between items. This additional information was essential to gauge participants’ progress in extracting temporal patterns, a key aspect of statistical learning that goes beyond mere segmentation. During retrieval practice (***Figure 3***), participants were presented with a picture from the previous image stream and were prompted to recall the subsequent image. The cue remained on screen for three seconds. Participants then saw four options: the accurate image, a luring image from the same sequence (within-sequence lure), a luring image from a different sequence within the same prior image presentation (cross-sequence lure), and a fourth option "none above" (correct when the cue image was the final image of a sequence). Participants were asked to make a choice within 3 seconds. Each retrieval practice entailed six questions, with the cue image randomly chosen from previously viewed image streams. No music was involved in this stage.

**Figure 3.**
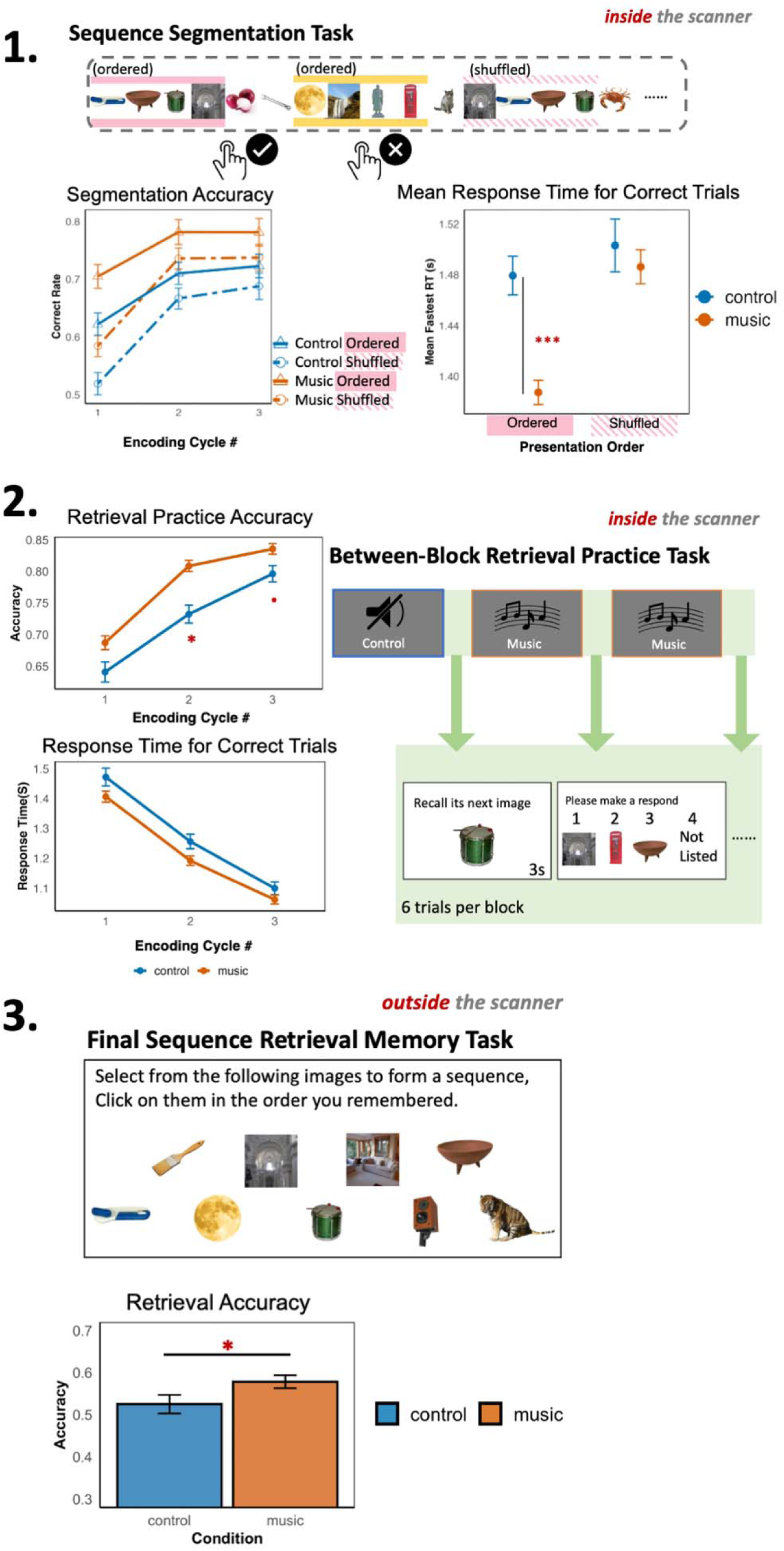
Learning Performance. **(1) *Sequence Segmentation Task:*** *Participants viewed continuous streams of images in ordered or shuffled sequences while pressed buttons to indicate perceived event boundaries. The segmentation accuracy plot shows higher performance in music conditions (orange) compared to control conditions (blue), particularly for ordered sequences (solid lines) versus shuffled sequences (dashed lines). Response times were faster for ordered sequences in the music condition. **(2) Between-Block Retrieval Practice:** During scanning, participants completed retrieval practice tasks after each learning block. Participants showed better accuracy and faster response times in the music condition (orange) compared to control (blue) across three encoding cycles. The task involved selecting the next image in a sequence from four options, with six trials per block. **(3) Final Sequence Retrieval:** After scanning, participants reconstructed complete sequences by selecting and ordering images from a pool of options. Performance was significantly better for sequences learned with musical accompaniment compared to those learned in silence*.

##### Final Retrieval Task

Following the encoding phase, participants exited the scanner and engaged in two memory retrieval tasks on a computer, each took around 3-5 minutes. The first was an item recognition task. Thirty items randomly selected from the encoding task and additional twenty new images were used. For each trial participants would see an image and they needed to decide if they have seen that image during the task. One purpose of this task was to build a time buffer to enable a test of delayed recall of the sequence memory (next task), and to provide an assessment of individual item familiarity which could help interpret performance levels in that more challenging recall task (i.e., if they don’t recognize the individual items well we can expect poor performance recalling the relational information). Then they performance a sequence retrieval task (***Figure 3***). Each of the 18 sequences from encoding was tested once. For each trial, participants viewed nine images on-screen for each question. Among these, four belonged to one sequence, while the remaining five were randomly selected from other sequences learned within the same run as the to-be-tested sequence. Participants had to select, group, and reorder these images into a familiar sequence by clicking on them sequentially.

Notably, no music or feedback was provided during retrieval practice or the final retrieval task, restricting interpretation of measured impacts of the music manipulations on memory to the encoding phase.

##### Practice Trial

Prior to the main encoding phase, participants completed a 5-minute practice trial to familiarize themselves with the task structure. The practice consisted of four sequences presented in a format identical to the main task: each sequence appeared six times with a 1/3 probability of random order shuffling. The practice sequences were presented without musical accompaniment to establish baseline task understanding. Following the practice sequences, participants completed a sequence reconstruction task where they grouped and ordered the images they had observed. This practice phase served two purposes: (1) to familiarize participants with the continuous stream presentation and segmentation response requirements, and (2) to demonstrate the task goal of identifying and learning sequential relationships among images. All practice stimuli were selected from the same image database as the main task but were not used in the experimental trials to prevent interference effects.

### A Statistical Learning Framework

Our paradigm implements core principles of statistical learning through several key design features. First, sequences appeared in their correct order only 2/3 of the time, with 1/3 probability of random shuffling. This probabilistic structure requires participants to extract statistical regularities rather than memorize deterministic associations—if a sequence A-B-C-D appeared 6 times per block, participants would experience it in correct order ∼4 times and shuffled ∼2 times. Learning therefore required computing distributional statistics across exposures to identify the underlying sequential structure. Second, sequences were embedded in continuous streams with variable numbers of lure images between them, preventing simple temporal counting strategies and requiring genuine pattern detection from noisy input. Third, the final memory test assessed generalization of learned statistical knowledge—participants had to reconstruct complete sequences from memory, demonstrating that they had extracted the underlying sequential relationships rather than just memorized specific stimulus co-occurrences.

This design extends our prior designs (Ren et al., 2024; Ren & Brown, 2025) by now capturing the essence of real-world statistical learning: extracting reliable patterns from variable, noisy experience where the signal (correct sequence) must be distinguished from noise (shuffled presentations and intervening lures). Critically, our paradigm differs from simple associative learning in several ways. Associative learning typically involves deterministic stimulus-response pairings that can be acquired through single-trial learning or straightforward repetition (e.g., A always follows B). In contrast, statistical learning requires accumulating evidence across multiple variable exposures to extract probabilistic relationships (e.g., A follows B 67% of the time). Our participants could not rely on simple associative mechanisms because: (1) sequences were probabilistically rather than deterministically structured, (2) lure stimuli created interference requiring pattern extraction from noise, and (3) successful performance required generalization to novel test conditions (reconstruction without environmental cues).

While we did not test generalization of this learning to entirely novel sequences, our paradigm assessed generalization across multiple dimensions: (1) from encoding context (with/without music) to retrieval context (always without music), (2) from probabilistic presentation during learning (2/3 correct) to deterministic reconstruction requirements during testing, and (3) from sequential presentation during encoding to simultaneous presentation during reconstruction. These forms of generalization demonstrate that participants extracted underlying statistical structure rather than memorizing surface-level stimulus co-occurrences.

### fMRI acquisition

Brain scans were collected using a 3T Siemens Prisma scanner with a 32-channel head coil at the GSU/GT Center for Advanced Brain Imaging. High-resolution T1-weighted structural images were obtained using magnetization prepared rapid gradient echo (MPRAGE) sequences with a repetition time (TR) of 2500 ms, echo time (TE) of 2.22 ms, and field of view (FOV) of 256 × 240 mm. The sequence used 0.8 mm slice thickness with 0.8 mm isotropic voxel size, an 8° flip angle, and 220 Hz/pixel bandwidth. GRAPPA parallel imaging was employed with an acceleration factor of 2.

T2-weighted structural images were acquired using 3D SPACE sequence with GRAPPA parallel imaging (acceleration factor = 2). The sequence parameters included a TR of 3200 ms, TE of 563 ms, and FOV of 256 × 240 mm. The images were acquired with 0.8 mm slice thickness, 0.8 mm isotropic voxel size, 120° flip angle, and 744 Hz/pixel bandwidth.

T2*-weighted functional data and localizer data were obtained using Multiband echo planar imaging (EPI) sequences with an acceleration factor of 5. The sequence employed a TR of 750 ms, TE of 32 ms, and FOV of 220 × 220 mm. Images were acquired with 2.5 mm isotropic voxel size across 72 slices, using a 52° flip angle and 2367 Hz/pixel bandwidth. The T2*-weighted images provided whole-brain coverage and were acquired with slices oriented parallel to the long axis of the hippocampus.

Detailed hippocampal scans were obtained in T2 space with selective excitation using ZOOMit 2D TSE (Turbo Spin Echo) provided by Siemens. The sequence parameters included a TR of 4560 ms, TE of 51 ms, 2 mm slice thickness, 130° flip angle, and 100 Hz/pixel bandwidth.

All participants were provided with foam ear plugs and a noise-cancelling MRI-compatible headset. This equipment served dual purposes: enabling communication with the experimenter between runs in the scanner and optimizing music stimuli delivery during scanning. Although all music stimuli were standardized to the same volume level, participants were asked to adjust the sound volume inside the scanner using an MRI-compatible controller before task initiation. They were instructed to set the volume to a level where the music was clearly audible in the MR environment while maintaining comfort and safety. Consequently, while the absolute intensity level of the sound varied across participants, the relative intensity remained consistent across all audio stimuli within each participant’s session.

### fMRI preprocessing

Results included in this manuscript come from preprocessing performed using fMRIPrep 23.2.1 (Esteban et al., 2019) (RRID:SCR_016216), which is based on Nipype 1.8.6 (Gorgolewski et al., 2011) (RRID:SCR_002502). We implemented standard preprocessing steps including motion correction, slice timing correction, spatial normalization to MNI space, and physiological noise regression. B0 field inhomogeneity correction was applied using topup. Full preprocessing details, including all parameters and implementation specifics, are provided in Supplementary Document.

### fMRI analysis

The fMRI analysis employed multiple complementary approaches to characterize how music influences sequence learning at both regional and network levels. We began with traditional univariate analyses using generalized linear models (GLM) to identify brain regions showing differential activation across task conditions. Given our specific hypotheses about the roles of memory-related regions, we conducted focused analyses within anatomically-defined regions of interest (ROIs) – medial temporal lobe (particularly the hippocampus and the parahippocampal cortex), striatum (caudate and NAc in particular) and ventromedial prefrontal cortex. To understand how these regions work together, we examined functional connectivity patterns during learning.

#### Univariate Generalized Linear Model

After preprocessing, a generalized linear regression model (GLM) analysis was performed to understand how functional BOLD responses were modulated by different task conditions. The GLM analysis utilized FSL (FMRIB Software Library, Version 6.0.5.6; Jenkinson et al., 2012), specifically FEAT (FMRI Expert Analysis Tool) for first- and higher-level analyses (Woolrich et al., 2004).

Before running the GLMs, the preprocessed data from fmrirep was further smoothed with a 5mm full-width at half maximum Gaussian kernel in FEAT. The model included six nuisance regressors accounting for head motion (translational and rotational), twelve possible image/task conditions during the visual stream presentation (context: music or control; sequence presentation order: ordered or shuffled; position of the image: within-sequence (image 1-3) or boundary (image 4)), and four conditions during retrieval practice (See ***Table 1***). The task regressors were convolved with a double-gamma hemodynamic response function, and temporal derivatives were included. The complete design matrix comprised 38 regressors per run.

**Table 1.** Regressor Labels and Conditions for GLM.

| Regressor Label | Sequence Learning |  |  |
| --- | --- | --- | --- |
|  | Context Condition | Sequence Order | Location of the Image |
| CO_WT | Silence/Control | Ordered | Within Event |
| CO_BD | Silence/Control | Ordered | Boundary |
| CS_WT | Silence/Control | Shuffled | Within Event |
| CS_BD | Silence/Control | Shuffled | Boundary |
| MO_WT | Music | Ordered | Within Event |
| MO_BD | Music | Ordered | Boundary |
| MS_WT | Music | Shuffled | Within Event |
| MS_BD | Music | Shuffled | Boundary |
| Lure_CO | Silence/Control | Ordered | Non-member Lure |
| Lure_CS | Silence/Control | Shuffled | Non-member Lure |
| Lure_MO | Music | Ordered | Non-member Lure |
| Lure_MS | Music | Shuffled | Non-member Lure |
|  | Retrieval Practice |  |  |
|  | Image Condition during Learning | Task Phase |  |
| Cue_C | Control | Cue shown |  |
| Ans_C | Control | Subjects make answer |  |
| Cue_M | Music | Cue shown |  |
| Ans_M | Music | Subjects make answer |  |

Group-level analyses were performed using FSL’s FLAME (FMRIB’s Local Analysis of Mixed Effects) stage 1 to account for both within-subject and between-subject variability. Final cluster results were thresholded using a cluster-corrected threshold of Z > 3.1, with a cluster significance threshold of p < 0.05.

#### Regions of Interest

Based on our theoretical framework and prior literature on sequence learning and schema effects, we identified three key neural circuits as regions of interest (ROIs):

1. Medial temporal lobe (MTL) regions: The hippocampus and parahippocampal cortex were selected based on extensive evidence for their role in binding sequential elements (Hsieh et al., 2014), detecting temporal regularities (Schapiro et al., 2014, 2016), and forming associative memories (Ranganath, 2010). The known functions of these regions are critical for the integration component of our Cross-Modal Temporal Scaffolding theory.
2. Prefrontal regions: The ventromedial prefrontal cortex (vmPFC) was selected based on its established role in schema-guided learning (van Kesteren et al., 2012), while the inferior frontal gyrus (IFG) was included due to its involvement in rule extraction and cognitive control during statistical learning (Karuza et al., 2013). Testing for modulation of these regions allows us to test for neural correlates of the functional balance in the CMTS framework between areas argued to be fundamental for schema integration (vmPFC) versus those known to be taxed by explicit control processes for learning (IFG); to the extent that music scaffolds learning, this balance and thus the neural pattern, should shift.
3. Striatal regions: The nucleus accumbens (NAc) and caudate were selected for their established roles in processing rhythmic patterns (Grahn & Rowe, 2009) and temporal predictions (Bornstein & Daw, 2012). These regions are particularly relevant given that musical temporal structure might influence temporal prediction mechanisms during statistical learning.
4. Angular gyrus: This region was included based on its role in integrating multimodal information and semantic processing (Seghier, 2013), which should be thus modulated by any facilitation of the higher-order integration processes in our task.

This selection of ROIs enables us to test both hypothesized mechanisms of the CMTS framework: (1) enhanced memory integration through MTL-vmPFC interactions, and (2) reduced demands on explicit control resources through shifts away from IFG-parietal networks.

The IFG, AG, parahippocampal gyrus (anterior and posterior), caudate, and NAc masks were extracted in FSL using the Harvard-Oxford Atlas (Desikan et al., 2006). The hippocampus masks were manually traced for each participant using ITK-SNAP (www.itksnap.org; Yushkevich et al., 2006) on high-resolution hippocampus scans using the finalized protocol developed by the Hippocampal Subfields Group (*in prep*; https://hippocampalsubfields.com/harmonized-protocol/). To align these traced masks with other neuroimaging data, we employed a two-step registration process. First, FSL’s FLIRT (FMRIB’s Linear Image Registration Tool) was used to calculate the transformation matrix between the high-resolution hippocampus scan and the T1 scan, allowing us to realign the traced masks into the T1 native space. Subsequently, these realigned masks were transformed into MNI (Montreal Neurological Institute) standard space using ANTs (Advanced Normalization Tools), ensuring spatial consistency across participants for group-level analyses.

For the vmPFC, considering its broad anatomical coverage, the masks were 5mm spheres centered on coordinates reported by the lab’s prior studies (anterior vmPFC: MNI = [-2, 56,-10]; posterior vmPFC: MNI = [-5, 16, -11], reported in (Brown et al., 2016)), which our ongoing work implicates in using schemas to support new learning.

#### Functional Connectivity

One aim of this study was to understand how different brain regions interactively contribute to statistical learning and sequential learning, and how the networks were modulated by music. To address this question, we performed a seed-based functional connectivity analysis and an ROI-to-ROI connectivity analysis (including graph theory) using CONN Toolbox (Whitfield-Gabrieli & Nieto-Castanon, 2012). Preprocessed data from fmriprep was imported in to CONN for further denoising to minimize the influence of physiological and motion-related confounds. This CONN’s default pipeline implemented the anatomical CompCor approach (Behzadi et al., 2007), which included: (1) regression of white matter and CSF signals using principal component analysis (5 components each), (2) regression of motion parameters and their first-order temporal derivatives, (3) identification and scrubbing of motion-affected time points (frame-wise displacement > 0.5 mm), (4) linear detrending, and (5) band-pass filtering (0.008-0.09 Hz). For seed-based gPPI analysis, we limited the seed ROIs to: hippocampus, amygdala, parahippocampal gyrus, caudate, NAc and vmPFC.

For our ROI to ROI connectivity analyses, we expanded beyond our primary ROIs to include additional regions implicated in sequence learning. While our initial ROI selection focused on regions with well-established roles in our theoretical framework, the connectivity analyses benefited from a more comprehensive network approach. We therefore included: 1) Core ROIs from our univariate analyses: Hippocampus, parahippocampal cortex (anterior and posterior), vmPFC, IFG, angular gyrus, NAc, and caudate. 2) Additional frontal regions: Superior frontal gyrus and middle frontal gyrus were included based on their consistent appearance in our univariate results from both current and previous study (Ren & Brown, 2025) and their documented involvement in executive functions supporting sequence/associative learning (Bischoff-Grethe et al., 2004; Strange et al., 2001). 3) Auditory cortex(superior temporal gyrus) was included to account the music’s effects on perceptive systems; 4) Additional subcortical and limbic regions: The amygdala was included given its strong interconnections with the hippocampus and its appearance in our univariate results especially at sequence boundaries. The putamen was included alongside the caudate to provide a more complete assessment of striatal contributions to sequence learning. This expanded ROI selection allowed us to more comprehensively characterize network reorganization during music-accompanied statistical learning. By including regions spanning frontal-parietal control networks, limbic/memory systems, and striatal prediction circuits, we could better test our hypothesis that musical context shifts processing from variable frontal-parietal engagement toward more consistent MTL-vmPFC-striatal connectivity.

The vmPFC masks were again derived from prior schema-related studies, using 5mm spheres. Other ROIs were created directly using CONN’s default AAL atlas (Tzourio-Mazoyer et al., 2002). For functional connectivity analysis, we focused on the contrast between control and music during the whole encoding period because this comparison would reveal how music systematically altered the coordination between memory encoding system and other distinct networks such reward systems, during sequence learning, potentially explaining music’s facilitatory effects on memory formation.

Finally, to examine the relationship between neural connectivity patterns (including their stability across individual participants) and learning outcomes while addressing the high-dimensional nature of connectivity data relative to sample size, we implemented a random forest machine learning approach. For each condition (music and control), we constructed a separate model using the randomForest package in R. The input features consisted of all pairwise ROI-to-ROI functional connectivity beta values extracted from the CONN toolbox analysis, with each connection representing a single predictor variable. The target variable was each participant’s final retrieval accuracy score. We used the caret package in R with random forest regression, employing 5-fold cross-validation to balance model stability with our sample size constraints (n = 35 participants). Models were configured with 100 trees and nodesize = 3 to prevent overfitting. The optimal number of features per tree split (mtry parameter) was determined through automated grid search across values of 2, 155, and 309. For each mtry value, complete 5-fold cross-validation was performed, with model performance assessed using cross-validated R². The mtry value yielding the highest average cross-validated performance was selected for the final model fitted on the complete dataset. To establish whether model performance exceeded chance levels, we conducted permutation testing with 1000 iterations. For each permutation, target accuracy scores were randomly shuffled while maintaining the connectivity feature structure, creating null distributions for comparison with observed cross-validated performance. P-values were calculated as the proportion of permuted models achieving R² values equal to or greater than the observed performance. Given the high-dimensional connectivity data relative to sample size, we focused on comparing relative performance between conditions and examining optimal model architectures (mtry values) as indicators of underlying network complexity rather than making absolute claims about prediction accuracy.

## Results

### Behavioral Results: Musical Context Enhances Statistical Learning Performance

To help quantify whether musical context signals facilitate temporal event segmentation, participants completed an event boundary detection task during image viewing. Participants were tasked with pressing a button whenever they perceived an image as the ‘end of an group’ or an ‘event boundary’. It’s important to note that participants were not informed about any association between the images and music, and they were explicitly instructed to focus solely on the images and their sequential relationships. Nonetheless, it was hypothesized that participants might benefit from the presence of the music backdrop in segmenting the images into groups. To assess this, the successful event boundary recognition rate was calculated at each event boundary. A successful detection was counted when a participant pressed a button(s) at the last image of the sequence or the luring image(s) following the last image but not pressing buttons during the middle of a sequence (both boundary detection and non-boundary rejection). Average detection accuracy (percentage of trials that the boundary was successfully detected) was calculated for each participant each run and each condition (context*presentation order). ***Figure 3.1*** displays the event boundary detection rates across three encoding runs. A repeated measures ANOVA was conducted testing run, context and presentation order as main effects showed significant effects of all three variables (run: F(2, 1007) = 28.674, p < .001, ηp² = .05; context: F(1, 1007) = 18.742, p < .001, ηp² = .02; presentation order: F(1, 1007) = 19.038, p < .001, ηp² = .02, see details in Supplementary Table 1). Post-hoc pairwise comparisons, using Bonferroni correction for multiple comparisons, revealed consistently higher boundary detection rate in music condition (music ordered > control ordered; music shuffled > control shuffled; see Supplementary Table 2 for detailed statistical significance). Beyond accuracy, we analyzed reaction times for correct boundary detections to further understand the efficiency of event segmentation processes. Reaction time was measured as the time from the onset of the boundary picture (the last picture in a sequence) to the participant’s first button press after this onset and before the start of the next sequence. A repeated measures ANOVA examining the effects of condition (music vs. control) and presentation order (ordered vs. shuffled) on reaction times revealed significant main effects of both factors and a significant interaction between them (condition: F(1, 20275) = 21.017, p < .001, ηp² = .001; presentation order: F(1, 20275) = 30.239, p < .001, ηp² = .002; interaction: F(1, 20275) = 6.218, p = .013, ηp² = .0003). As shown in ***Figure 3.1***, post-hoc comparisons revealed that the reaction time advantage for music was specific to ordered sequences. For ordered sequences, participants responded significantly faster during the music condition compared to the control condition (difference = 0.092 seconds, p < .001). However, for shuffled sequences, there was no significant difference in reaction times between music and control conditions (difference = 0.017 seconds, p = .493).

The retrieval practice tests inserted during encoding provided an opportunity to assess participants’ learning progress over time and provide crucial evidence that musical benefits extend beyond boundary detection. By calculating the correct retrieval rate for visual sequences in both the control and music conditions, an average learning curve was constructed (***Figure 3.2***). During the tests, no music was present, yet participants showed significantly enhanced accuracy for sequences originally learned with musical accompaniment. A repeated measure ANOVA treating subject as repeated measure testing run and context condition as main effect suggested significant effects of these two variables (run: F(2, 517) = 42.05, p < .001, ηp² = .14; context: F(1, 517) = 9.522, p = .002, ηp² = .02; see Supplementary Table 3). A post-hoc pairwise comparison using Bonferroni correction indicated a significant higher retrieval accuracy in music condition at run 2 and a trend effect at run 3 (run 2: mean difference = -0.071, 95% CI [-0.133, -0.009], p = .026; run 3: mean difference = -0.053, 95% CI [-0.116, 0.010], p = .099; see Supplementary Table 4 for details). In addition to accuracy, we analyzed reaction times for correctly answered trials in the retrieval practice task to assess processing efficiency. A repeated measures ANOVA with run and condition as factors revealed a significant main effect of condition (F(1, 515) = 4.362, p = .037,, ηp² = .008), with faster responses in the music condition compared to the control condition. Although post-hoc pairwise comparisons with Bonferroni correction did not reveal statistically significant differences for specific runs after adjusting for multiple comparisons, the overall pattern indicated faster processing for sequences learned with musical accompaniment (***Figure 3.2***).

After encoding, participants performed an item recognition task and a sequence retrieval task. Item recognition memory performance was analyzed using signal detection theory to account for both hits and false alarms. For sequence items learned in the music condition, participants showed high discrimination sensitivity (d’ = 3.95, SD = 0.88), with a hit rate of 96.4% (SD = 9.1%) and a false alarm rate of 4.9% (SD = 10.5%). Similarly robust performance was observed for sequence items from the control condition (d’ = 4.07, SD = 0.88; hits = 96.9%, SD = 9.8%; false alarms = 4.9%, SD = 10.5%). Non-sequence lure items that were not part of any temporal structure also showed comparable discrimination (d’ = 3.96, SD = 1.00; hits = 95.7%, SD = 10.7%; false alarms = 4.9%, SD = 10.5%).

A repeated measures ANOVA comparing discrimination sensitivity (*d’*) across the three item types revealed no significant differences between conditions in simple item recognition (F(2,222) = 0.092, p = .912), suggesting that the presence of musical context signals during encoding did not differentially affect subsequent item recognition. The consistently high *d’* values across all conditions indicate that participants maintained strong item recognition memory regardless of the learning context or temporal structure. Two participants were identified as outliers (exceeding 2 SD from the mean d’) and were excluded from subsequent analyses, leaving a total behavioral results sample size of 87.

By contrast, in the Final Retrieval Task, participants were required to identify images belonging to the same sequence/event but also to correctly order them. While statistical learning is often considered an implicit learning process, the data suggested the possibility of explicit learning of the sequential relationships between images. This was particularly noteworthy given that the sequential relationships were probabilistic rather than deterministic during encoding, with each sequence appearing in its fixed order only 66.6% of the time. For each sequence, participants received credit only if they reconstructed the entire sequence in its correct order (i.e., all images placed in their exact serial positions). Under these scoring criteria, participants successfully reconstructed 57.8% of sequences (SD = 30.6%) in music condition, while those sequences in the control condition were reconstructed with 52.5% correct rate (SD = 31.7%). Consistent with our hypothesis that musical context would enhance temporal sequence learning, a paired-samples t-test revealed significantly higher sequence reconstruction accuracy in the music condition compared to the control condition (t(86) = -2.292, p = .024, d = .246) (***Figure 3.3***).

### Whole-brain Univariate GLM Results

Building on the behavioral findings, to understand the whole-brain neural patterns during the statistical learning and temporal order memory task modulated by music, we conducted a univariate GLM analysis using FSL’s FEAT. Based on previous findings suggesting music might reduce neural demands during sequence learning (Ren & Brown, 2025), I first examined the contrast between music and control conditions for the entire encoding phase regressors. ***Figure 4*** shows the regions that exhibited different BOLD activity during the two conditions. Images and sequences learned without music (control) showed stronger BOLD activity in prefrontal regions including the superior frontal gyrus and middle frontal gyrus, as well as the lateral occipital cortex. In contrast, the music condition showed stronger BOLD activity in music and auditory-processing-related regions including the superior temporal gyrus and precentral gyrus (an expected outcome and useful sanity check, given the control condition in this study was sequencing in silence). Detailed activation clusters are provided in ***Table 2***.

**Figure 4.**
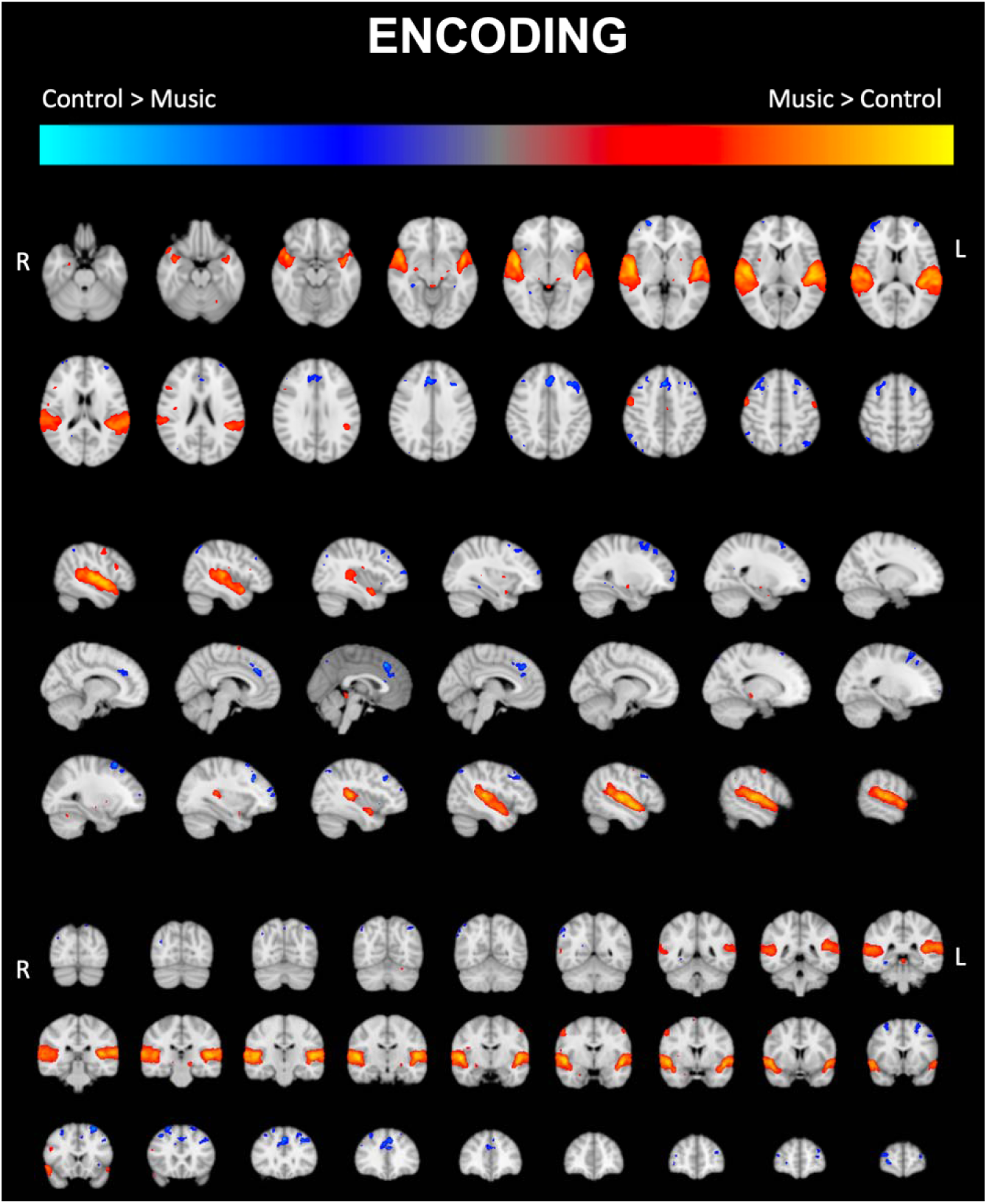
Whole-Brain Univariate GLM Results for Encoding Phase. *Regions with significant BOLD activity differences between music and control conditions during the encoding phase are displayed. Control showed stronger activation in prefrontal and occipital regions, while music activated auditory and motor-related areas*.

**Table 2.** Clustering Results for Whole-Brain GLM Contrast during Encoding.

| Cluster Index | Voxels | P | Z-MAX | Z-MAX X (mm) | Z-MAX Y (mm) | Z-MAX Z (mm) | Brain Region |
| --- | --- | --- | --- | --- | --- | --- | --- |
| **Control > Music** |  |  |  |  |  |  |  |
| 7 | 557 | 1.10E-09 | 5.62 | 2 | 28 | 42 | Paracingulate Gyrus |
| 6 | 391 | 1.79E-07 | 4.63 | 24 | 24 | 62 | Superior Frontal Gyrus [R] |
| 5 | 351 | 6.56E-07 | 4.6 | -34 | 26 | 36 | Middle Frontal Gyrus [L] |
| 4 | 260 | 1.62E-05 | 4.72 | -28 | 18 | 60 | Superior Frontal Gyrus [L] |
| 3 | 190 | 0.00025 | 4.22 | 44 | -62 | 56 | Lateral Occipital Cortex [R] |
| 2 | 178 | 0.000413 | 4.48 | 28 | 62 | 2 | Frontal Pole [R] |
| 1 | 77 | 0.05 | 4.08 | -44 | -68 | 50 | Lateral Occipital Cortex [L] |
| **Music > Control** |  |  |  |  |  |  |  |
| 3 | 5546 | 0 | 8.86 | 52 | -10 | 4 | Superior Temporal Gyrus [R] |
| 2 | 4805 | 3.08E-43 | 8.42 | -48 | -6 | -6 | Superior Temporal Gyrus [L] |
| 1 | 195 | 0.000204 | 5.12 | 54 | -2 | 50 | Precentral Gyrus [R] |

We then investigated the control vs. music contrast separately for the regressors specifically modeling the boundary (BD) vs. within the sequence (WT) to better understand how musical context affects different aspects of sequence processing. The activation maps are shown in ***Figure 5***, and the detailed cluster results are presented in ***Table 3*** and ***Table 4***. *At the boundary*, the control > music contrast again revealed stronger activity in frontal and occipital regions. Music, on the other hand, activated not only the expected auditory regions due to the nature of the stimulus, but now there was also a significant difference in the amygdala and thalamus. When examining activation when processing images *within the sequence*, the control condition showed stronger effects in the lingual gyrus and retrosplenial cortex (precuneus) in addition to prefrontal and occipital regions. The music condition continued to show stronger activity in auditory-related areas, including Heschl’s gyrus and precentral gyrus (***Figure 4***, bottom).

**Figure 5.**
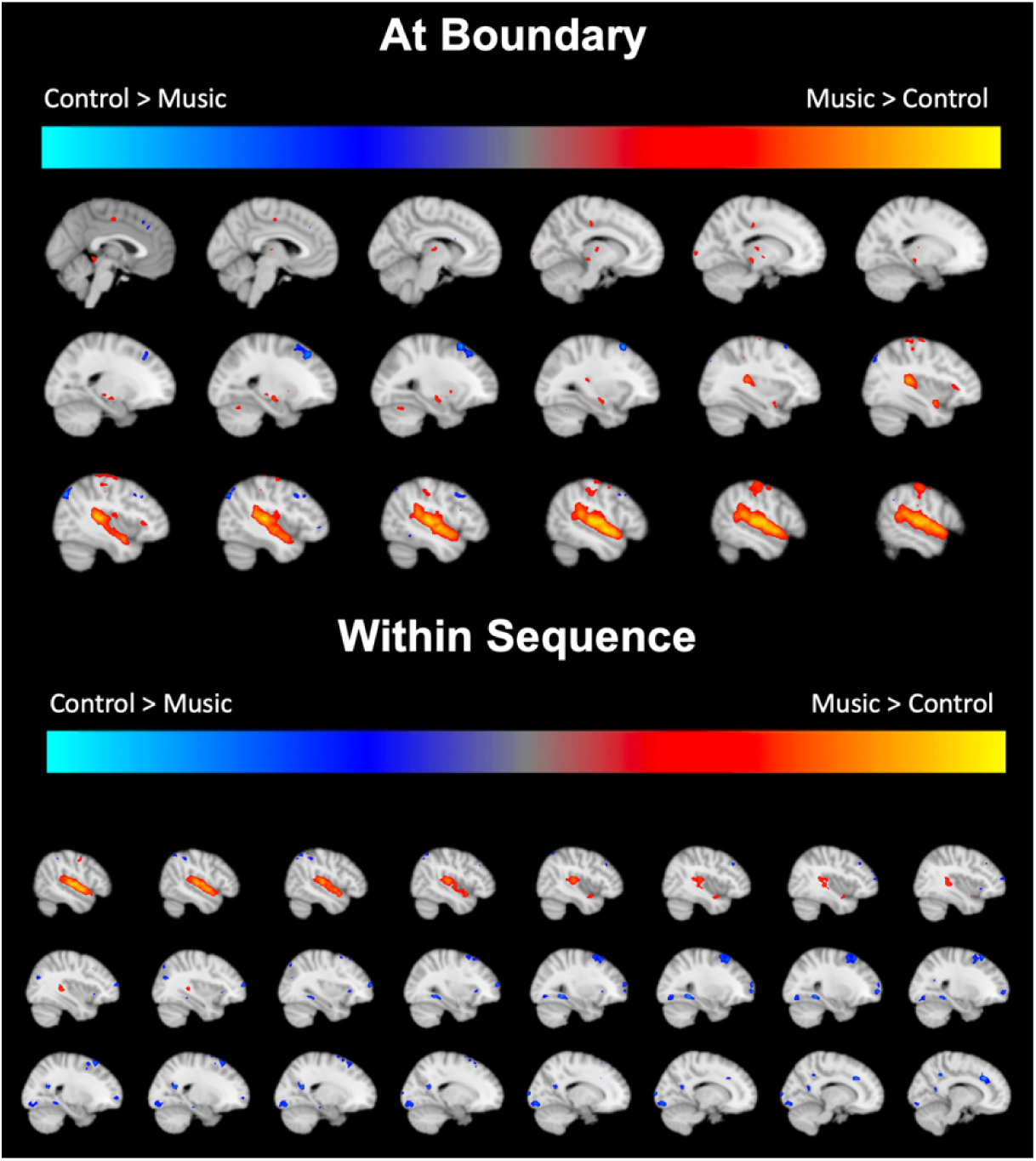
Univariate GLM Contrast – Music vs Control at Event Boundaries and within Sequence. *This figure shows the contrast between music and control conditions at sequence boundaries (top) and within sequence (bottom)*.

**Table 3.**
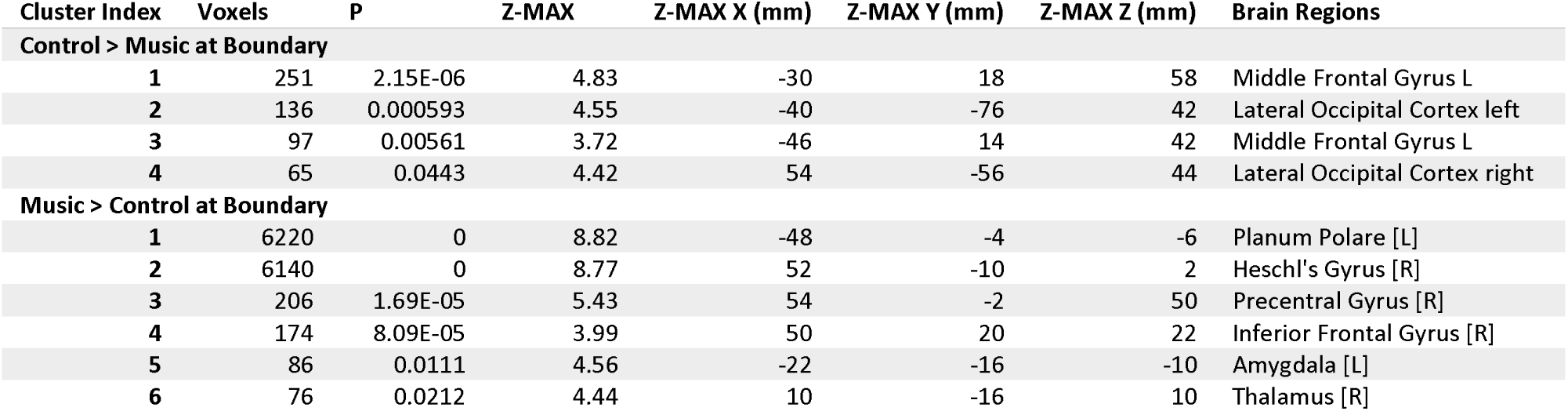
Clustering Results for GLM Contrast at Event Boundaries (Control vs. Music)

**Table 4.** Clustering Results for GLM Contrast Within Sequence (Control vs. Music)

| Cluster Index | Voxels | P | Z-MAX | Z-MAX X (mm) | Z-MAX Y (mm) | Z-MAX Z (mm) | Brain Region |
| --- | --- | --- | --- | --- | --- | --- | --- |
| **Control > Music within Sequence** |  |  |  |  |  |  |  |
| 1 | 888 | 2.18E-15 | 4.76 | -12 | -104 | 6 | Occipital Pole [L] |
| 2 | 568 | 3.35E-11 | 5.21 | 10 | 28 | 34 | Paracingulate Gyrus [R] |
| 3 | 357 | 5.96E-08 | 4.51 | 24 | 24 | 60 | Superior Frontal Gyrus [R] |
| 4 | 277 | 1.31E-06 | 4.1 | 16 | -92 | -8 | Occipital Pole [R] |
| 5 | 176 | 0.000117 | 4.44 | -28 | 58 | 10 | Frontal Pole [L] |
| 6 | 170 | 0.000156 | 4.6 | 34 | 60 | 12 | Frontal Pole [R] |
| 7 | 158 | 0.000282 | 4.55 | -34 | 26 | 36 | Middle Frontal Gyrus [L] |
| 8 | 156 | 0.000311 | 4.89 | -28 | -56 | -4 | Lingual Gyrus [L] |
| 9 | 141 | 0.000669 | 4.79 | 28 | -46 | -8 | Lingual Gyrus [R] |
| 10 | 123 | 0.00174 | 4.39 | 20 | -58 | 18 | Precuneous [R] |
| 11 | 89 | 0.0121 | 4.36 | 54 | -62 | 48 | Lateral Occipital Cortex [R] |
| **Music > Control within Sequence** |  |  |  |  |  |  |  |
| 1 | 3163 | 4.80E-37 | 8.38 | 52 | -12 | 4 | Heschl's Gyrus [R] |
| 2 | 2945 | 2.78E-35 | 7.82 | -50 | -22 | 8 | Heschl's Gyrus [L] |
| 3 | 158 | 0.000282 | 4.79 | 54 | -2 | 46 | Precentral Gyrus [R] |

Lastly, to investigate how encoding context influences subsequent retrieval processes, we conducted a contrast analysis during the retrieval practice period, focusing on the cue and response phases. As shown in ***Figure 6***, during the 3-second cue presentation, trials that were originally learned in silence showed stronger activity in the supramarginal gyrus in both hemispheres compared to those learned with music. During participants made responses for the retrieval practice task, the control condition exhibited stronger activity in the occipital pole and lateral occipital cortex. The decreased activation in the music condition, particularly in regions associated with visuospatial processing and attention (supramarginal gyrus, occipital cortex), may reflect lower cognitive effort required to retrieve temporal relationships when sequences were initially encoded with the structured temporal framework. This interpretation aligns with our behavioral findings showing enhanced sequence memory in the music condition, suggesting that musical context during encoding may facilitate the formation of more readily accessible temporal representations.

**Figure 6.**
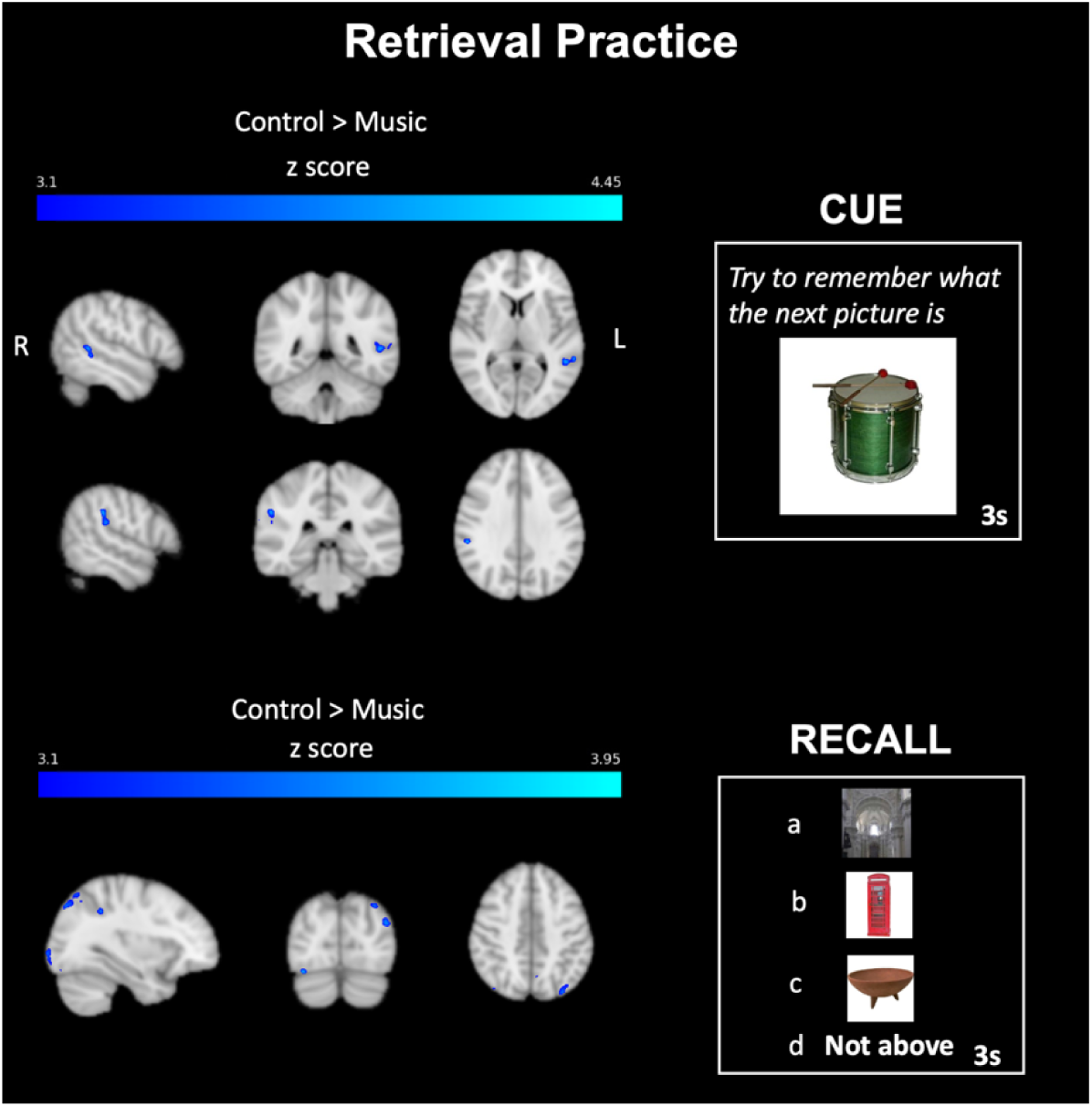
Retrieval Practice Period Univariate Contrast. *BOLD activity during retrieval practice is shown for both cue and response phases. The control condition activated the supramarginal gyrus and occipital cortex more strongly, highlighting distinct retrieval strategies between music and control conditions*.

In summary, these whole-brain analyses revealed distinct patterns of neural activation between music and control conditions during both encoding and retrieval, demonstrating the substantial influence of musical context on brain activity during statistical learning. The results partially supported our hypotheses: as predicted, we observed reduced neural demands in prefrontal and visual processing regions during music-accompanied learning. However, contrary to our predictions, we did not observe enhanced MTL activity during boundary detection in the music condition. Instead, music engaged subcortical regions (amygdala, thalamus) during boundary processing. The absence of significant MTL differences between conditions in the whole-brain analysis warrants careful interpretation. Given the MTL’s fundamental role in sequence learning and temporal order processing, this region likely remains critically engaged regardless of learning context. This consideration, combined with our specific hypotheses about MTL function in music-accompanied sequence learning, motivated a more focused ROI analysis. The following section examines targeted ROIs – theoretically relevant regions identified from prior literature and our previous work, allowing us to more precisely characterize how musical context modulates activity in these critical memory circuits.

### ROI-Behavior Correlations Reveal Efficient Learning with Music

Based on the extant literature and our theoretical framework of CMTS, we identified several key ROIs in which to test their relationship with statistical learning and its enhancements with music. Specifically, we focused on three neural circuits believed to be critical for sequence learning: (1) medial temporal lobe regions (hippocampus, anterior and posterior parahippocampal cortex) given their established role in binding sequential elements and detecting temporal regularities (Hsieh et al., 2014; Schapiro et al., 2014, 2016); (2) ventromedial prefrontal cortex (vmPFC), which interacts with the MTL in schema-guided learning (van Kesteren et al., 2012; Preston & Eichenbaum, 2013); and (3) striatal regions (nucleus accumbens and caudate), which process rhythmic patterns and predictions (Grahn & Rowe, 2009; Bornstein & Daw, 2012). We also included the inferior frontal gyrus (IFG) as a control region associated with explicit rule learning and cognitive control.

Before testing our predictions about network reconfigurations between these functional nodes through which music might benefit learning, we first examined the individual relationships between ROI activity during encoding and subsequent memory performance. We hypothesized that if an ROI’s functions for visual sequential statistical learning are facilitated processing in the presence of music, one would expect negative correlations between BOLD activity and performance, indicating more efficient neural processing.

We tested this prediction by correlating subject-level beta values and subjects’ subsequent retrieval test performance separately for music and control conditions. The results (***Figure 6***) revealed a striking pattern of negative correlations between neural activity and performance during encoding with music in context. In both the hippocampus and anterior PHC, as well as vmPFC and striatal regions (NAc and caudate), we observed significant negative correlations between BOLD activity and behavioral performance specifically during within-sequence processing in the music condition (hippocampus: r = -.34, p = .004; aPHC: r = -.24, p = .045; vmPFC: r = -.43, p < .001; NAc: r = -.45, p < .001; caudate: r = -.27, p = .022). In contrast, these regions showed either weak positive or non-significant correlations in the control condition.

To formally test whether the correlation patterns between context conditions (music vs. control) were significantly different, we conducted Fisher’s Z-tests. To control for multiple comparisons across our six a priori ROIs, we applied False Discovery Rate (FDR) correction. After correction, the correlations between retrieval performance and BOLD activity were significantly more negative in the music condition for hippocampus (z = -2.24, p-FDR = .045), posterior PHC (z = -2.17, p-FDR = .045), NAc (z = -2.34, p-FDR = .045), and caudate (z = -2.22, p-FDR = .045). The vmPFC showed a marginally significant effect after correction (z = -1.96, p-FDR = .059), while anterior PHC showed a non-significant trend in the same direction (z = -1.70, p-FDR = .089). This pattern of results, with four of six a priori ROIs showing significantly different brain-behavior relationships after correction for multiple comparisons, strongly supports our hypothesis that musical context specifically modulates neural efficiency during sequential processing.

To assess whether these correlations reflected region-specific effects rather than global patterns, we examined the inferior frontal gyrus (IFG), a region associated with cognitive control. Unlike memory and reward regions, IFG showed no significant correlations with performance in either condition (all p > .08). This dissociation suggests musical context selectively modulates memory circuits rather than producing global changes across all task-engaged regions. These results support a mechanism through which music influences sequence learning: more efficient within-sequence processing across MTL, prefrontal, and striatal circuits, as evidenced by the consistent negative correlations (***Figure 7***). The consistency of these negative correlations across hippocampus, vmPFC, and striatum is particularly meaningful, as these regions form a circuit implicated in both schema-guided learning and temporal prediction. The coordinated efficiency across this network suggests that musical context optimizes the entire circuit rather than just individual components. This pattern aligns with our hypothesis that music provides an organizational framework that facilitates the encoding of sequential relationships.

**Figure 7.**
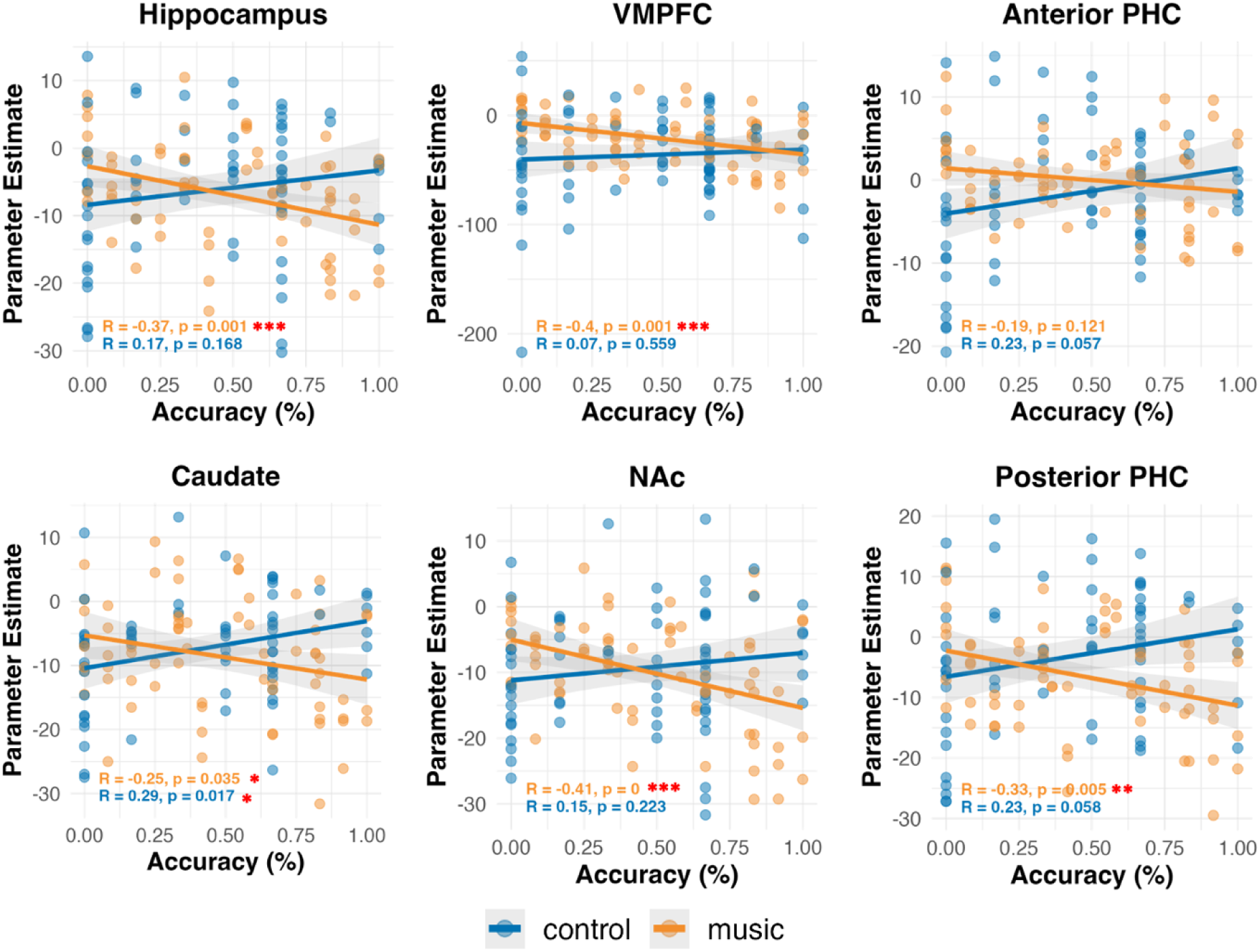
Correlation Between ROI Activity and Behavioral Performance. *This figure shows correlations between activity in ROIs (MTL, VMPFC, striatum) and subsequent retrieval performance. Each scatter plot shows individual participant data with regression lines indicating the direction and strength of correlations. The observed pattern reveals condition-specific relationships between neural activity and behavior: in the music condition (orange), several memory-related regions including the hippocampus, vmPFC, and striatum show significant negative correlations between activity during within-sequence (WT) processing and subsequent retrieval accuracy*.

### Seed-based gPPI and ROI-to-ROI Connectivity

The above data illustrate the distributed brain systems that are engaged differently for sequence learning under different contextual conditions in our study. To formally examine how musical context signals influence network-level interactions during sequence learning, we focused this study on generalized psychophysiological interaction (gPPI) analyses using the CONN toolbox and targeted seed regions of interest drawing on the prior work in our lab and others. Seed-based connectivity results are reported in ***Table 5*** and ***Table 6***, containing significant clusters found to be connected with each seed region differently for: 1) music vs. control overall; 2) music vs. control at event boundary; and 3) music vs. control within the sequence.

**Table 5.** Seed-Based Functional Connectivity Results for Music > Control.

| Music > Control |  |  |  |  |  |  |  |
| --- | --- | --- | --- | --- | --- | --- | --- |
| Cluster (x,y,z) | size | size p-FWE | size p-FDR | size p-unc | peak p-FWE | peak p-unc | regions |
| ENCODING OVERALL |  |  |  |  |  |  |  |
| **Amygdala** |  |  |  |  |  |  |  |
| 32 -74 +50 | 195 | 0 | 0 | 0 | 0.24061 | 0 | Lateral Occipital Cortex |
| 54 +30 +22 | 48 | 0.05188 | 0.01415 | 0.00659 | 0.99999 | 0.00008 | MidFG r, IFG |
| 52 +10 +36 | 41 | 0.10435 | 0.02196 | 0.00142 | 1 | 0.0001 | Precentral Gyrus |
| **Hippocampus** |  |  |  |  |  |  |  |
| -08 -54 54 | 70 | 0.00684 | 0.00524 | 0.00009 | 0.91482 | 0.00001 | Precuneous |
| **aPHC** |  |  |  |  |  |  |  |
| 52 -50 +36 | 83 | 0.00247 | 0.00194 | 0.00003 | 0.8281 | 0.00001 | IFG |
| 2 +30 +36 | 79 | 0.00346 | 0.00194 | 0.00005 | 0.98746 | 0.00002 | Retrosplenial Cortex |
| **pPHC** |  |  |  |  |  |  |  |
| 4 -88 4 | 117 | 0.00018 | 0.00013 | 0 | 0.99694 | 0.00003 | Lingual Gyrus, Occipital Pole |
| **Caudate** |  |  |  |  |  |  |  |
| 60 -28 12 | 46 | 0.06912 | 0.04191 | 0.00094 | 0.99943 | 0.00005 | Planum Temporale |
| -56 -28 10 | 42 | 0.1023 | 0.04191 | 0.00142 | 0.89618 | 0.00001 | Planum Temporale |
| AT BOUNDARY |  |  |  |  |  |  |  |
| **Hippocampus** |  |  |  |  |  |  |  |
| 30 22 56 | 88 | 0.00117 | 0.0006 | 0.00002 | 0.99481 | 0.00003 | SFG/SMA |
| 22 -88 -4 | 65 | 0.00983 | 0.00591 | 0.00013 | 0.9998 | 0.00005 | Fusiform Gyrus |
| -8 -88 -30 | 41 | 0.08877 | 0.03647 | 0.00114 | 0.99176 | 0.00002 | Cerebellum |
| **aPHC** |  |  |  |  |  |  |  |
| -4 -62 32 | 103 | 0.04961 | 0.04477 | 0.00022 | 0.87958 | 0.00001 | Precuneous |

**Table 6.**
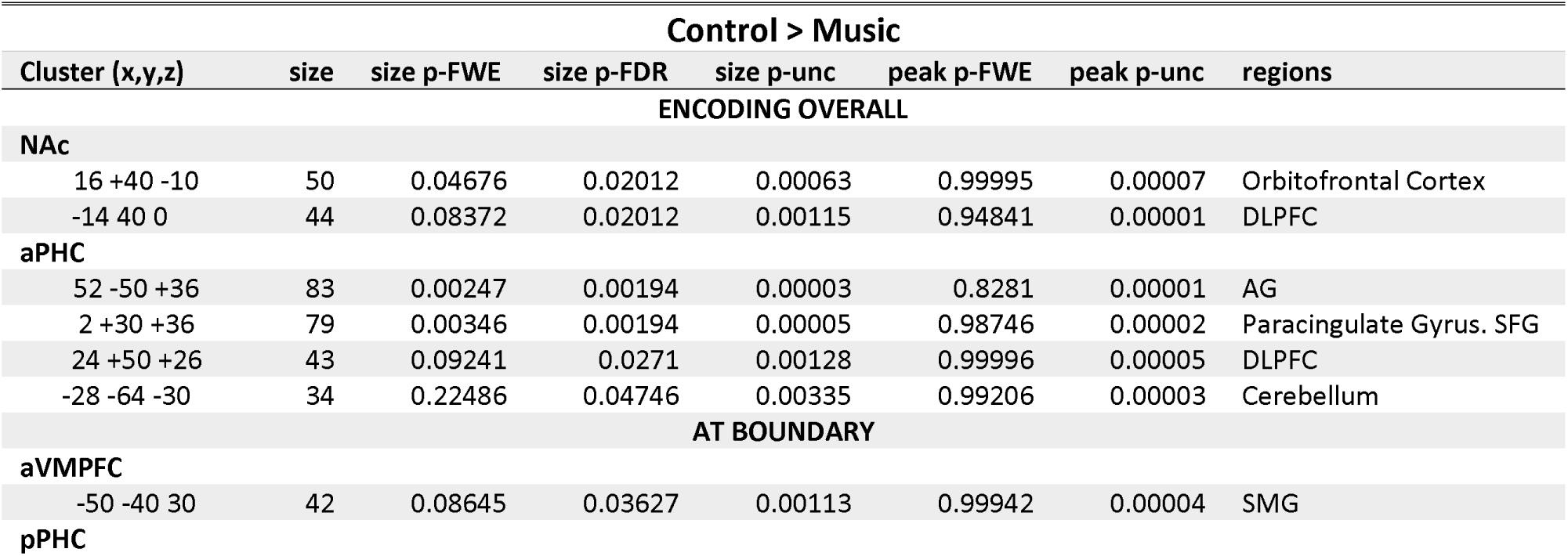
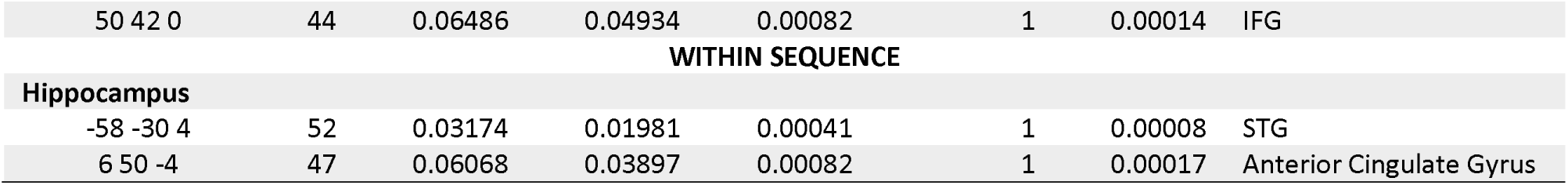
Seed-Based Functional Connectivity Results for Control > Music.

| Control > Music |  |  |  |  |  |  |  |
| --- | --- | --- | --- | --- | --- | --- | --- |
| Cluster (x,y,z) | size | size p-FWE | size p-FDR | size p-unc | peak p-FWE | peak p-unc | regions |
| ENCODING OVERALL |  |  |  |  |  |  |  |
| **NAC** |  |  |  |  |  |  |  |
| 16 +40 -10 | 50 | 0.04676 | 0.02012 | 0.00063 | 0.99995 | 0.00007 | Orbitofrontal Cortex |
| -14 40 0 | 44 | 0.08372 | 0.02012 | 0.00115 | 0.94841 | 0.00001 | DLPFC |
| **aPHC** |  |  |  |  |  |  |  |
| 52 -50 +36 | 83 | 0.00247 | 0.00194 | 0.00003 | 0.8281 | 0.00001 | AG |
| 2 +30 +36 | 79 | 0.00346 | 0.00194 | 0.00005 | 0.98746 | 0.00002 | Paracingulate Gyrus. SFG |
| 24 +50 +26 | 43 | 0.09241 | 0.0271 | 0.00128 | 0.99996 | 0.00005 | DLPFC |
| -28 -64 -30 | 34 | 0.22486 | 0.04746 | 0.00335 | 0.99206 | 0.00003 | Cerebellum |
| AT BOUNDARY |  |  |  |  |  |  |  |
| **aVMPFC** |  |  |  |  |  |  |  |
| -50 -40 30 | 42 | 0.08645 | 0.03627 | 0.00113 | 0.99942 | 0.00004 | SMG |
| **pPHC** |  |  |  |  |  |  |  |
| 50 42 0 | 44 | 0.06486 | 0.04934 | 0.00082 | 1 | 0.00014 | IFG |
| WITHIN SEQUENCE |  |  |  |  |  |  |  |
| Hippocampus |  |  |  |  |  |  |  |
| -58 -30 4 | 52 | 0.03174 | 0.01981 | 0.00041 | 1 | 0.00008 | STG |
| 6 50 -4 | 47 | 0.06068 | 0.03897 | 0.00082 | 1 | 0.00017 | Anterior Cingulate Gyrus |

The seed-based connectivity results (***Figure 8***, ***Table 5-6***) revealed a dissociation between music and control condition. The control condition demonstrated stronger connections between the medial temporal lobe (MTL) and striatum with prefrontal cortex regions including the dorsolateral prefrontal cortex (DLPFC), cingulate gyrus, and inferior frontal gyrus (IFG). This connectivity pattern aligns with our hypothesis that learning without musical structure would engage networks associated with explicit rule extraction and cognitive control (in our modeling analysis below, we show this is also a more idiosyncratic network profile). In contrast, in the music condition, MTL regions including the hippocampus, amygdala, and parahippocampus tended to show stronger connectivity with the visual information processing stream, including lateral occipital cortex, lingual gyrus, fusiform gyrus, as well as the precuneus, especially the retrosplenial cortex. This connectivity pattern between visual processing areas and MTL regions is consistent with our prediction that musical context would facilitate the active demands for integration of visual sequential information into memory circuits.

**Figure 8.**
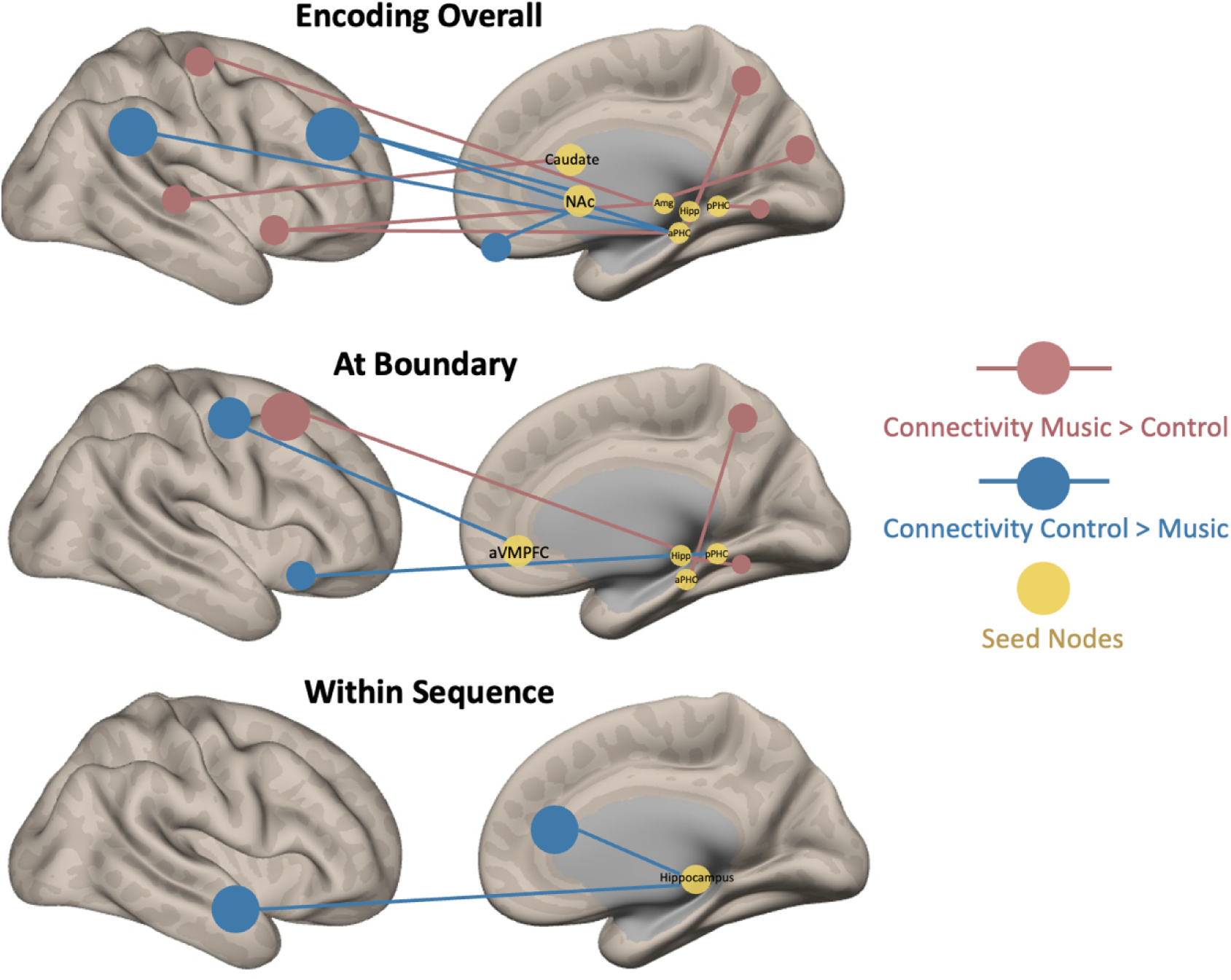
ROI-ROI Connectivity during Encoding. *Seed-based gPPI results showing differential connectivity patterns between music and control conditions across three analysis windows. (Top panel) Connectivity differences across the entire encoding phase; (Middle panel) Connectivity differences specifically at sequence boundaries; (Bottom panel) Connectivity differences within sequence processing. In all panels, yellow spheres indicate seed regions, red lines/regions represent significantly stronger connectivity in the music condition, and blue lines/regions represent significantly stronger connectivity in the control condition. The music condition predominantly enhanced connectivity between MTL regions and visual processing areas, while the control condition showed stronger connectivity between MTL/striatal regions and prefrontal cortex, supporting our hypothesis of distinct network engagement across conditions*.

The seed-based connectivity results provided initial evidence for distinct network reconfigurations across conditions. While informative, these results provided complex patterns of connectivity changes. To more systematically characterize network-level reorganization and directly test our theoretical predictions about specific circuit interactions, we conducted a comprehensive ROI-to-ROI connectivity analysis focusing on the key brain networks implicated in sequence learning.

The ROI-to-ROI connectivity analysis revealed two primary clusters showing significantly different connections between music and control conditions during sequence learning: one cluster (shown in red in ***Figure 9***) exhibited significantly higher connectivity during the music condition (F(2,33) = 8.45, p = .001, pFDR = .021). These clusters mainly connected the MTL and the ventromedial prefrontal cortex (vmPFC). For example, the posterior vmPFC and right hippocampus showed increased functional connectivity for the music condition during sequence encoding (t(34) = 3.3, p = .002, pFDR = .058). This enhanced MTL-vmPFC connectivity aligns with models of schema-guided learning and extends the seed-based observations of increased MTL integration with other brain regions during music-accompanied learning.

**Figure 9.**
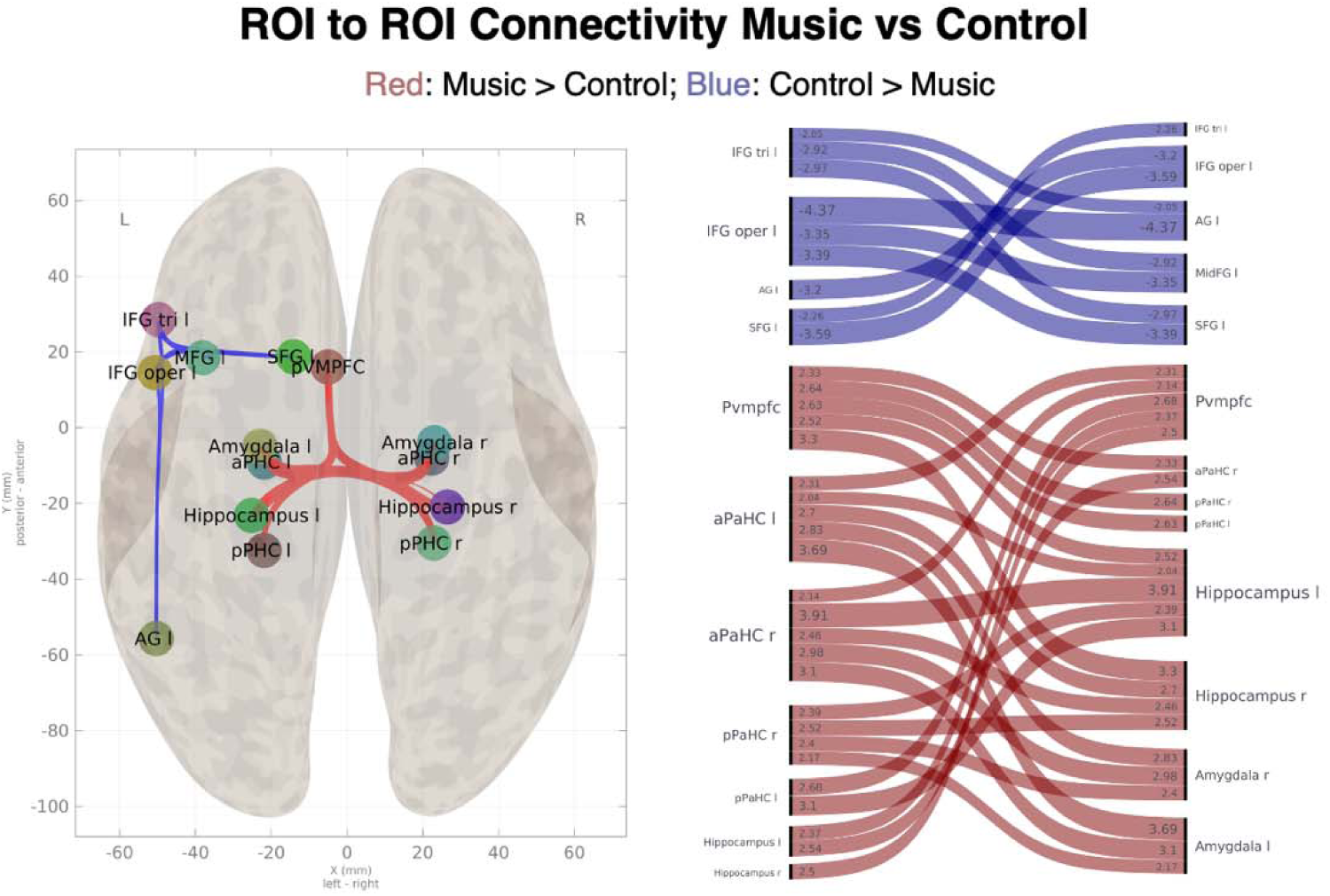
ROI-ROI Connectivity during Encoding. *This figure shows functional connectivity among ROIs for the music and control conditions. Music enhanced connectivity between the MTL and vmPFC. Control showed stronger IFG-parietal connectivity, linked to rule learning and syntactic processing*.

The second cluster (shown in blue in ***Figure 9***) demonstrated increased connectivity in the control condition (F(2,33) = 6.75, p = .003, pFDR = .035), predominantly located in the left hemisphere connecting the IFG with the inferior parietal lobule, a region highly associated with syntax and regularity learning, supporting our hypothesis that learning without musical structure would engage frontal-parietal control circuits..

### Network Connectivity Predicts Sequence Memory Performance in Music Condition

Having established that music fundamentally reorganizes functional brain networks during sequence learning, we next investigated whether these network configurations relate to behavioral outcomes. Specifically, we asked whether the distinct connectivity patterns observed in each condition could predict individual differences in learning success. Given the complexity of these network interactions and our relatively modest sample size, we employed a random forest modeling approach to identify which connectivity relationships most strongly predict memory performance in each condition.

Cross-validated random forest modeling revealed distinct connectivity-behavior relationships and network architectures between conditions. The music condition achieved a cross-validated R² of 0.387 with optimal mtry = 155, indicating that learning performance could be predicted from complex multi-network connectivity patterns. The control condition yielded R² of 0.32 with optimal mtry = 2, suggesting that participants’ performance on the task was explained by simpler, more focused connectivity patterns. Permutation testing (1000 iterations) revealed that the music condition showed a trend toward systematic connectivity-behavior relationships (p = 0.053 relative to chance, mean permuted R² = 0.242), while the control condition did not achieve reliable prediction above chance (p = 0.164, mean permuted R² = 0.209). The music condition demonstrated greater improvement over chance baseline (Δ = 0.145) compared to control (Δ = 0.111).

The optimal hyperparameters revealed fundamentally different network complexity requirements. The music condition’s optimal performance with mtry = 155 indicates that learning success required simultaneous integration across 155 connectivity features, suggesting complex multi-network coordination. In contrast, the control condition’s optimal performance with mtry = 2 indicates that learning could be predicted from simple patterns involving only 2 connectivity features at a time, suggesting more focused, local network relationships.

Variable importance analysis, examining which network interactions underpinned prediction of the model, revealed fundamentally different connectivity architectures. The control condition showed concentrated importance in local circuits in frontal and striatal areas, with putamen-NAc connection showing the highest importance, followed by IFG-MFG and other within-striatal connections. This pattern reflects focused, local network processing. In contrast, the music condition demonstrated distributed importance across diverse brain systems, including MTL-PFC interactions (caudate-vmPFC, IFG-hippocampus), limbic connections (SFG-parahippocampus), and PFC-striatal circuits (see supplementary document).

Taken together the network analysis findings collectively suggest that music modulates functional connectivity patterns during sequence learning, particularly enhancing connections between the MTL and visual processing regions, as well as between the MTL and vmPFC as well as between the PFC and the striatum. Furthermore, our modeling shows that the music condition also facilitates a more predictive relationship between brain connectivity and memory performance, highlighting the potential role of music in shaping neural networks involved in sequence learning and memory formation. These findings expand our understanding of how musical context shapes not just regional activity but also the broader neural networks supporting temporal sequence learning.

## Discussion

In this study, we used a novel statistical learning paradigm and investigated how musical context signals influence the neural mechanisms underlying visual sequence learning. Musical context consistently enhanced participants’ ability to detect event boundaries, improved sequence recall during interim tests, and led to better reconstruction of complete sequences. Using fMRI, we found this behavioral enhancement accompanied distinct changes in brain network organization and efficiency: music-accompanied learning showed reduced activation in fronto-parietal regions alongside more consistent engagement of memory-related networks especially between the medial temporal lobe and the prefrontal cortex, suggesting a shift from effortful sequence extraction to more efficient learning guided by familiar musical temporal structure. These findings reveal how structured auditory context can fundamentally alter the brain networks supporting visual sequence learning.

Our results demonstrate neural reorganization during statistical learning when structured temporal information from one modality influences sequence processing in another – a phenomenon we term "cross-modal temporal scaffolding" (***Figure 1***). This theoretical framework explains how musical context can facilitate (or in other cases - discussed briefly below – impair) visual statistical learning through two complementary mechanisms: enhanced memory integration through schema-guided learning, and reduced demands on explicit control resources.

### Memory Integration through Cross-Modal Temporal Scaffolding

“Schema theory” propose that when new information maps onto existing knowledge structures, learning becomes more efficient and robust (van Kesteren et al., 2012). These mental frameworks (“schemas”) facilitate rapid integration of incoming information by providing organizational scaffolding that reduces encoding demands. Music science has long utilized this same terminology to emphasize that music compositions can be highly schematic, and it is the adherence of compositions to learned syntactic structures that allow music to be processed with incredible efficiency (Leman, 2012). We argue that this schematic property enables music to exert predictable influences cross-modally on other learning processes. Musical context signals can vary in their syntactical “strength” (how closely they adhere to learned music structures; e.g., Western Classical), as well as veridical memory strength (how well the specific composition is known) – we have previously explicitly shown that both factors combine to upregulate parallel visual sequencing processes (Ren et al 2024; Ren & Brown, 2025). The Cross-Modal Temporal Scaffolding (CMTS) framework argues that, mechanistically, this benefit should (1) manifest from upregulated schema-guided learning mechanisms in the brain, reorganizing how the brain processes sequential information in the visual domain, and should (2) be highly impactful in statistical learning processes, because detecting the temporal regularities of *when* stimuli recur relative to one-another is a challenging mnemonic integration problem when that timing is variable.

In support of this, we observed a striking dissociation between networks engaged during statistical sequential learning without musical context versus with music accompaniment. Learning in silence predominantly engaged frontal-parietal connectivity centered on IFG-angular gyrus interactions – circuits associated with explicit rule extraction and grammatical learning (Karuza et al., 2013; Kim et al., 2022; Skosnik et al., 2002). This observation fits with our prediction that participants need to rely more heavily on effortful pattern extraction in the absence of an aid like musical scaffolding, and thus tax associated neural systems more strongly. In contrast, with music accompaniment, the same processes shifted toward distributed networks anchored by MTL-vmPFC connectivity, aligning with models of schema-mediated learning, where existing knowledge structures facilitate the integration of new information through dynamic cross-talk between these systems (Gilboa & Marlatte, 2017; van Kesteren et al., 2012). This result replicates and extends our previous finding of increased MTL-PFC connectivity during explicit deterministic visual sequence learning with music (Ren & Brown, 2025), suggesting that musical structure provides a robust scaffold for sequence encoding across different learning paradigms. Extending schema theory, CMTS proposes how such brain interactions can be triggered by purely temporal (rather than semantic) congruence across difference sensory domains.

An important test is whether music drives fundamentally different network architectures that translate to distinct learning mechanisms. Cross-validation random forest modeling revealed that the music condition achieved optimal model performance by integrating 155 connectivity features, with distributed importance across MTL-PFC-striatal circuits including caudate-vmPFC, IFG-hippocampus, and limbic-cortical connections. In contrast, the control condition model’s optimal performance involved only 2 connectivity features, suggesting focused and local network relationships. This architectural difference suggests different processing strategies, rather than control regions being ‘uninvolved’ in statistical learning. The control condition’s concentrated architecture aligns with explicit pattern extraction under uncertainty, engaging cognitive control circuits that show greater individual variability (Siegelman & Frost, 2015; Wang et al., 2017).

The network coordination in music condition reflects well-established anatomical and functional relationships that optimize learning through several mechanisms. The prefrontal cortex receives prediction-related feedback from the striatum and can modulate limbic system activity, including hippocampal encoding (Haber & Knutson, 2010), while the striatum and hippocampus can collaborate to integrate temporal and spatial contextual information during memory formation (Polti et al., 2022) as well as sequential retrieval (Brown et al., 2012; Brown & Stern, 2014)– even when relationships are learned with uncertainty or implicitly (Harrison et al., 2006; Schendan et al., 2003). The striatum’s established role in temporal prediction and rhythm processing (Grahn & Rowe, 2013) has been argued to be important for leveraging the temporal structure provided by music stimuli to generate predictions about upcoming sequence elements, while its interactions with PFC align with reward prediction error circuit that precipitate memory and decision making (Bialleck et al., 2011; Frank et al., 2019). In our task, ‘rewards’ are implicit and can be conceptualized as successful predictions of upcoming sequence elements, akin to prediction error operations in many real-world scenarios when an explicit reward isn’t involved. Central to CMTS, musical context provides a predictable backdrop against which visual sequence predictions become more salient—temporal transitions in music amplify order relationships between visual stream items by providing parallel predictive cues, offering additional evidence for sequence transitions when visual cues alone might be ambiguous. The MTL’s established capacity for binding arbitrary sequential elements into coherent memory representations (Ranganath & Hsieh, 2016) enhanced through connectivity with vmPFC, creates a pathway through which musical "schemas" facilitate rapid integration of new visual information into sequences. This interaction in our data aligns with both schema theory (van Kesteren et al., 2010) and complementary learning systems theory (McClelland et al., 2020), showing that musical structure serves as a previously-learned associative framework that interleaves with and enhances new visual sequence learning. Musical schemas create a rich contextual environment that optimally coordinates multiple learning systems, supporting emerging models that suggest successful learning depends on synchronized interaction between prediction, attention, and memory networks (Bassett & Mattar, 2017; Mattar et al., 2018). his complex architecture reflects the coordinated multi-system integration predicted by Cross-Modal Temporal Scaffolding Theory, and our data demonstrate that this enhanced pattern of interconnectivity is precisely what music facilitates during visual statistical learning.

Our findings suggest musical context optimizes this flexibility by providing a structured temporal framework bridging auditory and visual modalities. Familiar melodies offer predictable temporal patterns with syntactical structure that listeners understand, helping track time and event order. This shared temporal structure coordinates otherwise distinct neural mechanisms for sequence processing, creating a common reference system that aligns processing across brain networks. This explanation is supported by work showing that consistent network states predict better learning outcomes(Mattar et al., 2018) and that external structure can reduce individual variability in learning strategies (Armstrong et al., 2017).

Overall, these network interactions advance our understanding of cross-modal statistical learning beyond existing literature. While previous research has shown that statistical learning can operate across modalities (Conway & Christiansen, 2006; Mitchel & Weiss, 2011) and that signals in one perceptual modality can facilitate learning in another (Cunillera et al., 2010; Mitchel et al., 2014; Seitz et al., 2007; Thiessen, 2010), our results reveal specific neural systems through which temporal structure in one modality enhances learning in another. The network reorganization we observed—from variable frontal-parietal engagement in the control condition to consistent MTL-PFC-striatal connectivity with music—suggests that structured input in one modality creates more optimal neural states for processing sequential information generally, extending beyond modality-specific effects. These findings contribute to broader theories about how contextual factors shape learning network efficiency, demonstrating that using a second modality to provide contextual structure can fundamentally alter how the brain processes sequential information.

### Neural Efficiency: Reduced Demands on Frontal Regions

While our network analyses demonstrated enhanced memory integration through schema-guided processing, a complementary mechanism emerged in our whole-brain activation patterns: musical context also altered the efficiency of neural processing during statistical learning. We define neural efficiency as the combination of reduced activation with maintained or improved performance—a pattern indicating more optimized resource allocation rather than disengagement).

Whole-brain analyses revealed a distinct pattern of neural efficiency in the music condition, characterized by decreased activity in frontal, parietal and occipital regions during encoding and between-block retrieval practice. For example, we observed reduced activation in superior and middle frontal gyrus, paracingulate gyrus, and frontal pole - regions consistently implicated in rule learning, cognitive control and working memory maintenance (Bischoff-Grethe et al., 2004; Destrebecqz et al., 2005; Strange et al., 2001). These frontal-parietal regions typically show enhanced activation during statistical pattern extraction (Turk-Browne et al., 2009) and temporal order processing (Knutson et al., 2004).

Critically, these neural efficiency patterns correlated with participants’ behavioral responses. When learning with musical context, participants showed faster response times during both boundary detection and retrieval practice tasks while maintaining higher accuracy. This pattern parallels broader principles of neural efficiency where external support or scaffolding can reduce demands on specific brain networks - for instance, when prior knowledge reduces medial temporal lobe demands during episodic memory encoding (van Kesteren et al., 2012). While such streamlined neural processing typically emerges after years of specialized training—as seen in musicians processing speech (Patel, 2011) or chess experts recognizing objects (Bilalić et al., 2012) – our findings suggest music provides a form of "borrowed expertise"—immediately optimizing neural efficiency without requiring extensive training in visual sequence learning itself.

This neural efficiency signature persisted beyond encoding into the retrieval phase, even though no music was present during retrieval. During between-block retrieval practice, participants who had learned sequences in silence showed stronger activation in the supramarginal gyrus and occipital regions, areas associated with visual working memory (Grill-Spector et al., 2001) as well as sequence and statistical learning (Guidali et al., 2019). This heightened activation suggests greater reliance on explicit maintenance and manipulation of visual sequence information when learning occurred without musical structure. The reduced engagement of these regions in the music condition indicates that familiar musical structure during preceding encoding decreased demands on cognitive control resources—effects that persist through both learning and subsequent retrieval phases.

These findings challenge traditional models of cognitive load that would predict interference when adding auditory processing demands during visual learning. While our prior work has shown such disruption can occur when music lacks schematic properties (Ren et al., 2024), the present study demonstrates that familiar, syntactically-regular music qualitatively transforms how cognitive resources are allocated during learning. This contradicts simple models of attentional capacity that would predict division of resources leading to performance decrements (Navon & Gopher, 1979). Instead, our findings align with Baddeley’s foundational work on the partial segregation of modalities in working memory buffers—visual sequence information occupies the visuospatial sketchpad while musical information is processed in the phonological loop, allowing parallel processing without direct competition (Baddeley, 2020; RepovŠ & Baddeley, 2006). The CMTS framework builds on this insight, further suggesting that appropriately structured cross-modal input enhances (not just reduces interference) processing efficiency through complementary engagement of neural systems, particularly when the secondary modality provides temporal cues that align with natural segmentation processes. This observation and our CMTS framework align with emerging perspectives suggesting that contextual factors can *redistribute* rather than simply increase or decrease cognitive load (Sweller, 2020), creating a qualitative shift in how neural resources are allocated during learning. This may explain why musical context reduced rather than increased cognitive demands, with a key prediction being that these benefits should diminish as interference between modalities increases (e.g., in Stroop-like tasks where different information types compete within the same modality).

### Efficient Sequence Learning in MTL-PFC-striatal Network

Beyond regional activation differences, these efficiency effects extend to network-level interactions. We observed reduced functional connectivity between the inferior frontal gyrus and angular gyrus during music-accompanied learning. This circuit has been consistently implicated in explicit rule extraction under conditions of uncertainty (Karuza et al., 2013; Kim et al., 2022), with stronger engagement typically reflecting increased demands on explicit pattern detection. The reduced engagement of this circuit with musical context supports our interpretation that music reduces reliance on effortful control processes during statistical learning, shifting the burden from explicit rule-learning systems to more efficient memory integration processes.

To better understand how these activation differences relate to learning success, we examined the relationship between regional brain activity and behavioral performance. Multiple regions including the MTL, striatum, and vmPFC showed robust negative correlations with sequence learning success specifically when learning occurred with musical context. The consistency of these negative correlations suggests that musical context creates a more efficient processing state where reduced neural activity reflects more optimal encoding, aligning with findings in expertise research where reduced neural activation often reflects more efficient processing (Kelly & Garavan, 2005). While we cannot definitively claim these regions are engaging in schema-guided processing based solely on these correlations, the specificity of this pattern to the music condition, combined with the established roles of these regions in relational memory and temporal prediction (van Kesteren et al., 2012; Grahn & Rowe, 2013), is consistent with our theoretical framework. The coordinated efficiency across this network suggests that musical context optimizes the entire circuit rather than just individual components.

Critically, this efficiency effect supports the second mechanism of our CMTS framework: musical structure reduces demands on explicit cognitive control resources by providing an organizing framework that streamlines processing through stable facilitative network organization. The negative correlations between neural activity and performance indicate that the increased efficiency in neural engagement isn’t merely a sensory-bound consequence of musical presence on brain signals, but rather a functionally relevant reorganization that directly contributes to enhanced learning outcomes. This pattern is consistent with our theory’s prediction that temporal scaffolding drives more optimal engagement of learning networks, shifting processing from resource-intensive explicit control systems to more automatic, schema-guided integration processes. The present study focused on familiar, syntactically-valid music. Importantly, the CMTS framework also predicts when disruptive rather than facilitative effects would occur. When the temporal scaffold is weakened—for instance, with unfamiliar or unpredictable music (Ren et al., 2024)—we would expect impairment rather than enhancement of learning. Under such conditions, the beneficial shift from top-down control networks to MTL-mPFC interactions would likely be reversed or disrupted. In fact, this prediction of CMTS is supported by our other work (Ren & Brown, 2025), which demonstrates that unpredictable musical context indeed decreases MTL-PFC and striatum-MTL connectivity—the opposite pattern of what we observed with familiar, predictable music for statistical learning in the current study. Instead of reducing cognitive load, unpredictable cross-modal input should increase demands on top-down control for statistical learning, given the added difficulty of aligning (noisy) temporal statistics between both modalities. This mechanistic neural framework would explain the disruptive behavioral effects observed even in deterministic sequencing problems when using unpredictable musical accompaniment (Ren et al., 2024), and highlights the critical importance of a temporal scaffold’s qualities in determining whether cross-modal information facilitates or interferes with learning. On top of that, we hypothesized other limiting factors or constraints that may lead to diminished or reversed CMTS effect include 1) the temporal characteristics of the scaffold does not align with the optimal segmentation rate of the to-be-learned sequence; significant mismatches in temporal pacing between modalities would likely disrupt rather than facilitate learning. 2) CMTS benefits should diminish as the working memory load within each modality approaches capacity limits—if either the auditory scaffold or visual sequence becomes too complex, the cross-modal benefit would likely disappear as resources within each modality become overtaxed. 3) the effectiveness of temporal scaffolding likely varies with individual differences in modality-specific processing abilities and prior experience; those with musical training might show stronger CMTS effects due to enhanced processing of musical temporal structure. Testing these limiting factors and operational constraints will be crucial for validating and refining the CMTS framework in future research.

### Implications and Limitations

Our findings suggest new principles for understanding how contextual factors optimize learning networks. While all learning conditions in our task are ostensibly MTL-dependent, the shift from more variable frontal-parietal engagement to more consistent MTL-PFC-striatal connectivity with music scaffolding demonstrates that structured temporal context can fundamentally alter the neural network organization behind task performance. Drawing on these and prior data, the CMTS framework generates several testable predictions for future applied contexts: temporal scaffolding effects should be particularly pronounced in populations with reduced cognitive control resources, and structured auditory context should enhance learning of temporal sequences (such as language patterns or mathematical procedures) more than non-temporal material. Additionally, the framework predicts that individuals with variable or inefficient neural processing might show greater benefits from musical scaffolding interventions. These hypotheses require direct empirical testing across clinical and educational populations to validate the translational potential of cross-modal temporal scaffolding principles.

Several alternative explanations for music’s enhancement effects warrant consideration. First, musical context might simply increase general arousal or attention (Ünal et al., 2013), enhancing memory indirectly rather than through specific schema mechanisms. However, our findings showed benefits specifically for sequential relationships rather than individual items, and the network reorganization centered on MTL-vmPFC-striatal circuits known for temporal processing rather than arousal or emotional systems. The random forest model architecture difference provides compelling evidence for mechanistic reorganization beyond simple contextual effects.

A related concern is whether our effects simply reflect boundary cueing—that consistent music-sequence pairings allowed participants to use musical changes as cues for sequence boundaries rather than demonstrating true temporal scaffolding. However, several lines of evidence argue against this interpretation. First, our task battery assessed complementary measures not limiting to boundary detection. For example, musical benefits were measured during retrieval when music was absent (thus, no perceptual cues were present), indicating that familiar melodies enhanced the encoding of within-sequence relationships rather than merely facilitating segmentation through active perception of temporal markers. Importantly, our companion investigation of hippocampal sequence representations in this dataset (Ren et al., under review) also reveal that musical context enhances hippocampal representation for item relationship within sequences—changes occurring beyond boundary detection that directly correlates with memory performance. These findings demonstrate that musical scaffolding fundamentally alters how sequential relationships are encoded in hippocampal memory circuits, extending well beyond the attention or segmentation effects expected from simple boundary cueing.

Several design considerations highlight important future directions. While our silence control prioritized ecological validity, systematic manipulation of specific musical features could identify which elements are most crucial for optimizing learning circuits. The CMTS framework predicts that temporal scaffolding may generalize beyond music to any structured temporal information from one modality that provides organizational support for learning in another—for example, learned rhythmic visual cues scaffolding auditory learning, or tactile patterns enhancing visual sequence encoding. Testing these cross-modal combinations would establish whether our observed network reorganization represents a general principle of multi-modal learning or is specific to music-visual interactions. Additionally, individual differences in musical training, working memory capacity, and baseline statistical learning abilities likely modulate susceptibility to temporal scaffolding effects, warranting systematic investigation across developmental and clinical populations.

## Conclusion

In summary, our findings reveal how musical context fundamentally reshapes brain networks during statistical learning, providing mechanistic insights into the relationship between temporal contextual structures and functional brain organization. The Cross-Modal Temporal Scaffolding framework we propose explains how structured auditory input drives a shift from variable frontal-parietal engagement to more consistent MTL-PFC-striatal connectivity, optimizing learning through dual mechanisms of enhanced memory integration and reduced demands on cognitive control resources. This network reorganization has significant implications for understanding how environmental contexts shape learning capacity and efficiency. The network architecture differences provide direct evidence for CMTS dual mechanisms. Musical scaffolding creates rich, coordinated network states that optimally bind sequential information across multiple systems. The control condition’s concentrated reliance on striatal-frontal circuits reflects increased demands on explicit control resources typically required for statistical learning without temporal scaffolding. Beyond theoretical advances, these findings suggest new approaches for optimizing learning in educational settings and potential compensatory strategies for clinical populations with memory impairments or learning difficulties. Future research should explore when CMTS effects would and would not occur and test whether similar principles extend beyond music to other forms of structured temporal context.

