## Supplementary for "Music Scaffolds Visual Statistical Sequence Learning Through Network-Level Reorganization in the Brain"

**S Table 1** **ANOVA Results: Segmentation Performance Affected by Context, Presentation Order and Encoding Run**

| Effect | Df | Sum Sq | Mean Sq | F value | Pr(>F) |
| --- | --- | --- | --- | --- | --- |
| Run | 2 | 3.34 | 1.6723 | 28.674 | 7.74E-13*** |
| Context Condition | 1 | 1.09 | 1.093 | 18.742 | 1.65E-05*** |
| Presentation Order | 1 | 1.11 | 1.1104 | 19.038 | 1.41E-05*** |
| Run:Context | 2 | 0.26 | 0.1323 | 2.269 | 0.104 |
| Run:Order | 2 | 0.02 | 0.0088 | 0.15 | 0.86 |
| Context:Order | 1 | 0.01 | 0.005 | 0.086 | 0.769 |
| Run:Context:Order | 2 | 0 | 0.0016 | 0.027 | 0.974 |
| Residuals | 1007 | 58.73 | 0.0583 |  |  |

**S Table 2** **Post-hoc Pairwise Comparisons using Bonferroni’s Correction on Segmentation Performance**

| Run# | contrast | estimate | SE | df | t.ratio | p.value |
| --- | --- | --- | --- | --- | --- | --- |
| 1 | Control Ordered - Music Ordered | -0.1197 | 0.037 | 1007 | -3.232 | 0.0076*** |
| 1 | Control Ordered - Control Shuffled | 0.0653 | 0.037 | 1007 | 1.763 | 0.4693 |
| 1 | Control Ordered - Music Shuffled | -0.0369 | 0.037 | 1007 | -0.996 | 1 |
| 1 | Music Ordered - Control Shuffled | 0.185 | 0.037 | 1007 | 4.995 | <.0001*** |
| 1 | Music Ordered - Music Shuffled | 0.0829 | 0.037 | 1007 | 2.237 | 0.1532 |
| 1 | Control Shuffled - Music Shuffled | -0.1022 | 0.037 | 1007 | -2.759 | 0.0355* |
| 2 | Control Ordered - Music Ordered | -0.0451 | 0.037 | 1007 | -1.217 | 1 |
| 2 | Control Ordered - Control Shuffled | 0.0691 | 0.037 | 1007 | 1.865 | 0.3752 |
| 2 | Control Ordered - Music Shuffled | 0.0244 | 0.037 | 1007 | 0.66 | 1 |
| 2 | Music Ordered - Control Shuffled | 0.1141 | 0.037 | 1007 | 3.081 | 0.0127* |
| 2 | Music Ordered - Music Shuffled | 0.0695 | 0.037 | 1007 | 1.877 | 0.3652 |
| 2 | Control Shuffled - Music Shuffled | -0.0446 | 0.037 | 1007 | -1.205 | 1 |
| 3 | Control Ordered - Music Ordered | -0.0449 | 0.037 | 1007 | -1.213 | 1 |
| 3 | Control Ordered - Control Shuffled | 0.0503 | 0.037 | 1007 | 1.357 | 1 |
| 3 | Control Ordered - Music Shuffled | 0.014 | 0.037 | 1007 | 0.378 | 1 |
| 3 | Music Ordered - Control Shuffled | 0.0952 | 0.037 | 1007 | 2.57 | 0.0619~ |
| 3 | Music Ordered - Music Shuffled | 0.0589 | 0.037 | 1007 | 1.591 | 0.6721 |
| 3 | Control Shuffled - Music Shuffled | -0.0363 | 0.037 | 1007 | -0.979 | 1 |

**S Table 3** **ANOVA Results: Retrieval Practice Accuracy Affected by Context and Encoding Run**

| Effect | Df | Sum Sq | Mean Sq | F value | Pr(>F) |
| --- | --- | --- | --- | --- | --- |
| run | 2 | 3.697 | 1.849 | 42.05 | < 2e-16*** |
| context | 1 | 0.419 | 0.4186 | 9.522 | 0.00214*** |
| run:context | 2 | 0.014 | 0.0072 | 0.024 | 0.84837 |
| Residuals | 517 | 22.73 | 0.044 |  |  |

**S Table 4** **Post-hoc Pairwise Comparisons using Bonferroni’s Correction on Retrieval Practice Accuracy**

| run | contrast | estimate | SE | df | t.ratio | p.value |
| --- | --- | --- | --- | --- | --- | --- |
| 1 | control - music | -0.0459 | 0.0316 | 517 | -1.453 | 0.1468 |
| 2 | control - music | -0.0708 | 0.0316 | 517 | -2.239 | 0.0256** |
| 3 | control - music | -0.0528 | 0.032 | 517 | -1.652 | 0.0992 |

**fMRI Preprocessing**

### Preprocessing of B0 inhomogeneity mappings

A total of 1 fieldmaps were found available within the input BIDS structure for this particular subject. A B0-nonuniformity map (or fieldmap) was estimated based on two (or more) echo-planar imaging (EPI) references with topup (Andersson et al., 2003).

### Anatomical data preprocessing

A total of 1 T1-weighted (T1w) images were found within the input BIDS dataset. The T1w image was corrected for intensity non-uniformity (INU) with N4BiasFieldCorrection (Tustison et al., 2010), distributed with ANTs 2.5.0 (Avants et al., 2008, RRID:SCR_004757), and used as T1w-reference throughout the workflow. The T1w-reference was then skull-stripped with a Nipype implementation of the antsBrainExtraction.sh workflow (from ANTs), using OASIS30ANTs as target template. Brain tissue segmentation of cerebrospinal fluid (CSF), white-matter (WM) and gray-matter (GM) was performed on the brain-extracted T1w using fast (FSL (version unknown), RRID:SCR_002823, (Zhang et al., 2001)). Brain surfaces were reconstructed using recon-all (FreeSurfer 7.3.2, RRID:SCR_001847, Dale, Fischl, and Sereno 1999), and the brain mask estimated previously was refined with a custom variation of the method to reconcile ANTs-derived and FreeSurfer-derived segmentations of the cortical gray-matter of Mindboggle (RRID:SCR_002438, (Klein et al., 2017). A T2-weighted image was used to improve pial surface refinement. Brain surfaces were reconstructed using recon-all (FreeSurfer 7.3.2, RRID:SCR_001847, (Dale et al., 1999), and the brain mask estimated previously was refined with a custom variation of the method to reconcile ANTs-derived and FreeSurfer-derived segmentations of the cortical gray-matter of Mindboggle (RRID:SCR_002438, Klein et al. 2017). Volume-based spatial normalization to two standard spaces (MNI152NLin6Asym, MNI152NLin2009cAsym) was performed through nonlinear registration with antsRegistration (ANTs 2.5.0), using brain-extracted versions of both T1w reference and the T1w template. The following templates were were selected for spatial normalization and accessed with TemplateFlow (23.1.0, Ciric et al., 2022): FSL’s MNI ICBM 152 non-linear 6th Generation Asymmetric Average Brain Stereotaxic Registration Model [Evans et al., 2012 RRID:SCR_002823; TemplateFlow ID: MNI152NLin6Asym], ICBM 152 Nonlinear Asymmetrical template version 2009c [Fonov et al. (2009), RRID:SCR_008796; TemplateFlow ID: MNI152NLin2009cAsym].

### Functional data preprocessing

For each of the 4 BOLD runs found per subject (across all tasks and sessions), the following preprocessing was performed. First, a reference volume was generated, using a custom methodology of fMRIPrep, for use in head motion correction. Head-motion parameters with respect to the BOLD reference (transformation matrices, and six corresponding rotation and translation parameters) are estimated before any spatiotemporal filtering using mcflirt (FSL , Jenkinson et al., 2012). The estimated fieldmap was then aligned with rigid-registration to the target EPI (echo-planar imaging) reference run. The field coefficients were mapped on to the reference EPI using the transform. The BOLD reference was then co-registered to the T1w reference using bbregister (FreeSurfer) which implements boundary-based registration (Greve & Fischl, 2009). Co-registration was configured with six degrees of freedom. Several confounding time-series were calculated based on the preprocessed BOLD: framewise displacement (FD), DVARS and three region-wise global signals. FD was computed using two formulations following Power (absolute sum of relative motions,Power et al., 2014) and Jenkinson (relative root mean square displacement between affines, Jenkinson et al. (2002)). FD and DVARS are calculated for each functional run, both using their implementations in Nipype (following the definitions by (Power et al., 2014). The three global signals are extracted within the CSF, the WM, and the whole-brain masks. Additionally, a set of physiological regressors were extracted to allow for component-based noise correction (CompCor, Behzadi et al., 2007). Principal components are estimated after high-pass filtering the preprocessed BOLD time-series (using a discrete cosine filter with 128s cut-off) for the two CompCor variants: temporal (tCompCor) and anatomical (aCompCor). tCompCor components are then calculated from the top 2% variable voxels within the brain mask. For aCompCor, three probabilistic masks (CSF, WM and combined CSF+WM) are generated in anatomical space. The implementation differs from that of Behzadi et al. in that instead of eroding the masks by 2 pixels on BOLD space, a mask of pixels that likely contain a volume fraction of GM is subtracted from the aCompCor masks. This mask is obtained by dilating a GM mask extracted from the FreeSurfer’s aseg segmentation, and it ensures components are not extracted from voxels containing a minimal fraction of GM. Finally, these masks are resampled into BOLD space and binarized by thresholding at 0.99 (as in the original implementation). Components are also calculated separately within the WM and CSF masks. For each CompCor decomposition, the k components with the largest singular values are retained, such that the retained components’ time series are sufficient to explain 50 percent of variance across the nuisance mask (CSF, WM, combined, or temporal). The remaining components are dropped from consideration. The head-motion estimates calculated in the correction step were also placed within the corresponding confounds file. The confound time series derived from head motion estimates and global signals were expanded with the inclusion of temporal derivatives and quadratic terms for each (Satterthwaite et al., 2013). Frames that exceeded a threshold of 0.5 mm FD or 1.5 standardized DVARS were annotated as motion outliers. Additional nuisance timeseries are calculated by means of principal components analysis of the signal found within a thin band (crown) of voxels around the edge of the brain, as proposed by (Patriat et al., 2017). All resamplings can be performed with a single interpolation step by composing all the pertinent transformations (i.e. head-motion transform matrices, susceptibility distortion correction when available, and co-registrations to anatomical and output spaces). Gridded (volumetric) resamplings were performed using nitransforms, configured with cubic B-spline interpolation.

Copyright Waiver

The above boilerplate text in this [fMRI processing] section was automatically generated by fMRIPrep with the express intention that users should copy and paste this text into their manuscripts unchanged. It is released under the CC0 license.

### **Differential BOLD Activity Across Conditions**

While the whole-brain analyses provided valuable insights into broad activation patterns, we conducted targeted ROI analyses to test specific hypotheses about how musical context influences key memory circuits. These analyses offered greater statistical sensitivity than whole-brain comparisons by focusing on theoretically motivated regions. The BOLD activity within key regions of interest (ROIs) was extracted from the GLM model and further analyzed to understand how neural responses varied across context (control vs. music), presentation order (ordered vs. shuffled), and image position (boundary vs. within-sequence). A repeated measures ANOVA was used to address this question, and revealed that these regions were modulated differently by these factors. S.***Table 5*** summarizes the ANOVA results.

**S Table 5** **ANOVA Results for ROIs’ BOLD Activity Across Conditions**

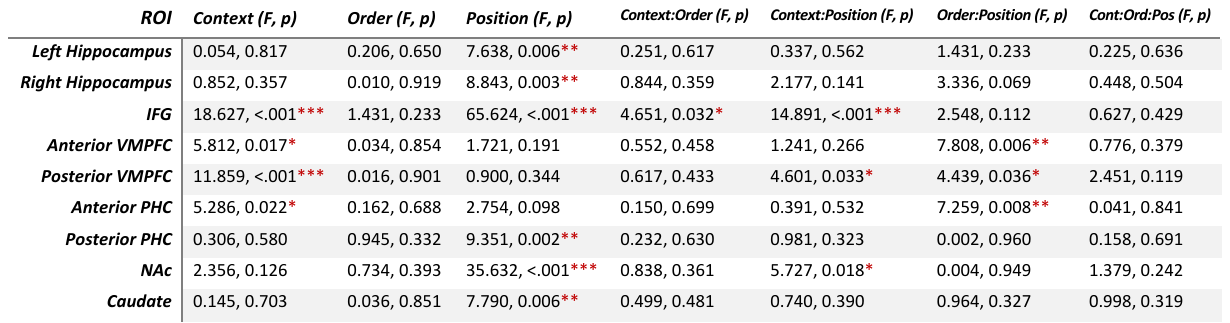

The hippocampus exhibited a significant main effect of position (left: F(1, 238) = 7.638, p = .006; right: F(1, 238) = 8.843, p = .003), with elevated activity for boundary images compared to within-sequence images (SFigure1). This boundary-specific enhancement patterns were also observed in posterior parahippocampal cortex (pPHC), nucleus accumbens (NAc), and caudate, suggesting their role in event boundaries processing.

The ventromedial prefrontal cortex (vmPFC) and anterior PHC showed demonstrated both a main effect of context and an interaction for order and position (aVMPFC: F(1, 238) = 7.808, p = .006; aPHC: F(1, 238) = 7.259, p = .008). As shown in SFigure1, both regions exhibited higher activity in the music condition than in the control condition. Moreover, both regions showed increased activity for sequence boundaries when presented in shuffled order with music. This pattern suggests these regions may be especially important for integrating temporal context when sequential relationships are less predictable.

Lastly, the inferior frontal gyrus (IFG) was significantly modulated by context (F(1, 238) = 18.627, p < .001) and image position (F(1, 238) = 65.624, p < .001), with an interaction effect between these factors (F(1, 238) = 14.891, p < .001). It showed stronger activity in the music condition compared to the control condition, as well as stronger activity for boundary than within-sequence images, especially in the music context. This aligns with my hypothesis of IFG’s role on how musical boundary affects visual boundary. Subsequent sections will continue to investigate this hypothesis.

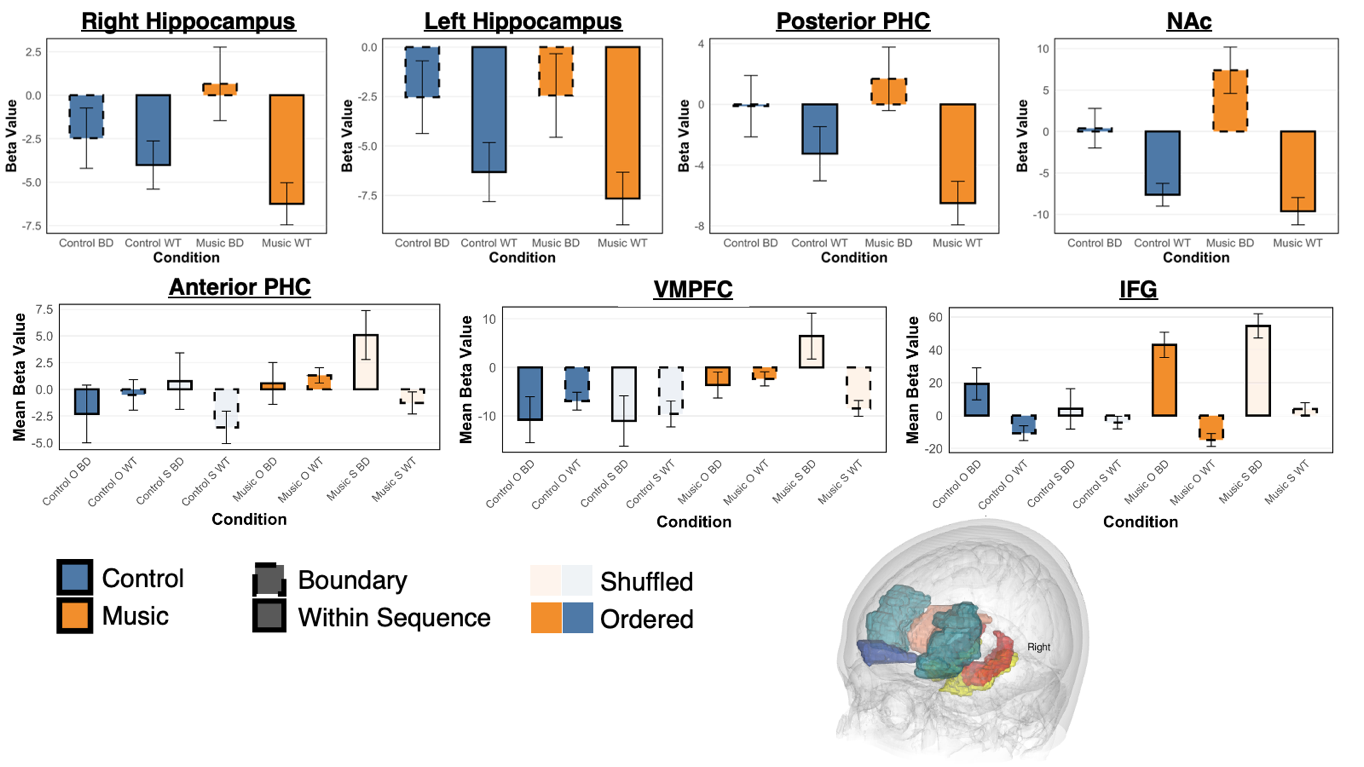

**S.Figure 1** **Differential BOLD Activity across Conditions for ROIs**

This figure illustrates BOLD activity for regions of interest. It emphasizes the modulation of activity by context (music vs. control), position (within sequence vs. boundary) and presentation order (ordered vs. shuffled).

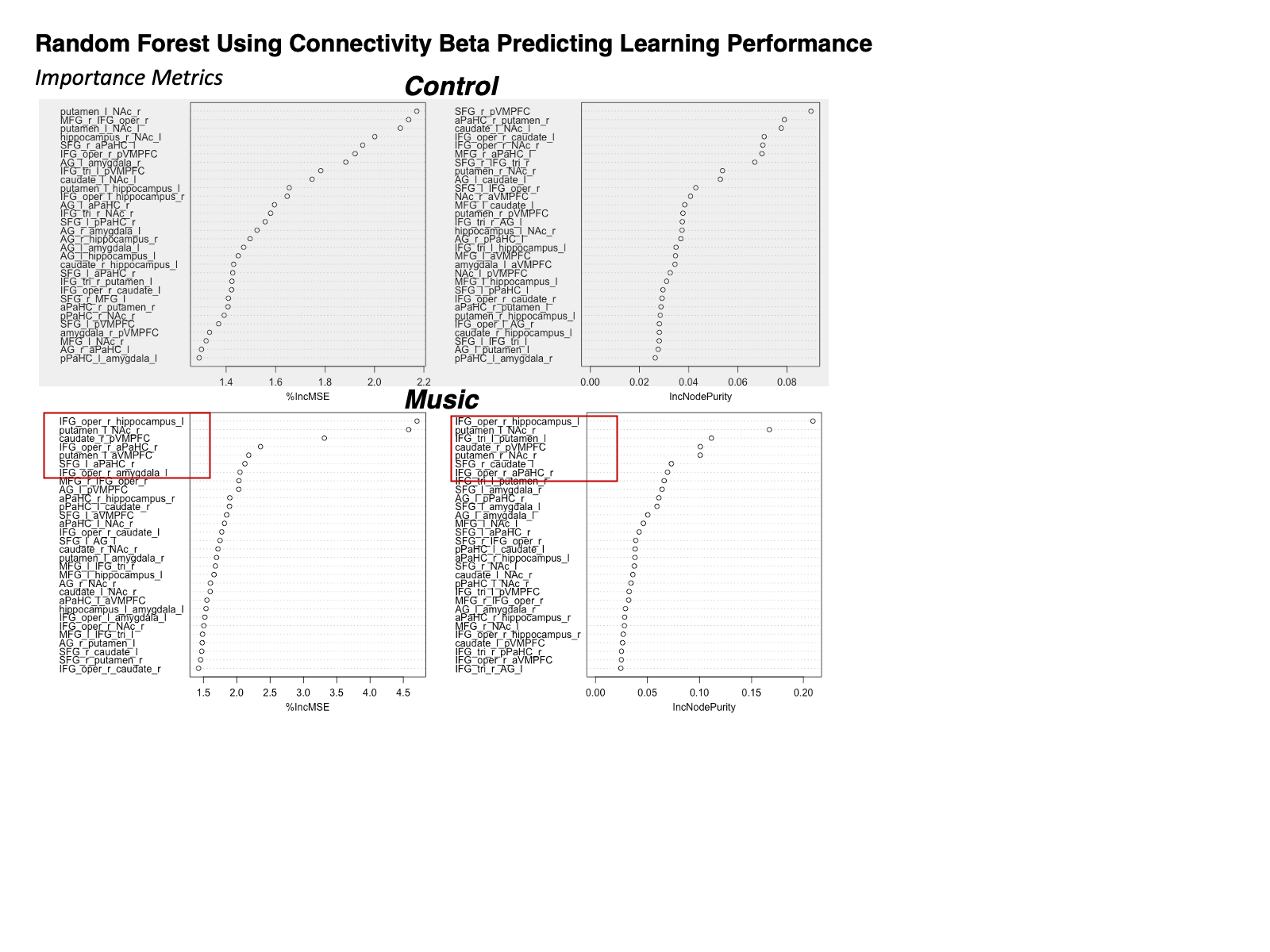
